# MyoGen: Unified Biophysical Modeling of Human Neuromotor Activity and Resulting Signals

**DOI:** 10.64898/2026.01.01.697284

**Authors:** Raul C. Sîmpetru, Ricardo G. Molinari, Devon R. Rohlf, Rebeka L. Batichotti, Renato N. Watanabe, Leonardo A. Elias, Alessandro Del Vecchio

## Abstract

Understanding human motor control requires integrating cortical and spinal cord activity, muscle mechanics, and electrophysiology, levels that are often studied separately. We present MyoGen, an open-source framework that unifies spinal circuitry, proprioceptive feedback, musculotendon dynamics, cortical activity, and multimodal electromyography (EMG) generation in a single, interoperable platform. Spinal motor neurons and the resulting motor unit (MU) action potentials represent the only neural cells that can be accessed at scale in humans. We used human MU ensembles to validate our model across a wide range of experimental conditions. Using data-driven optimization, we found that MyoGen produces MU population activity that closely matches experimental discharge-rate distributions and discharge variability across human muscles. Moreover, it generates decomposable surface and intramuscular EMG, reproduces beta-band modulation of descending drive and its nonlinear transformation into force, and implements complete sensorimotor loops. Dimensionality reduction of simulated agonist–antagonist EMG reveals low-dimensional control manifolds consistent with experimental recordings from both healthy and spinal cord injured individuals. MyoGen provides physiologically grounded, ground-truth data and integrates seamlessly with analysis pipelines, enabling systematic investigation of motor control principles, validation of signal-processing algorithms, and exploration of sensorimotor interactions that are experimentally inaccessible.

## 1 Introduction

Wearable neural interfaces are currently undergoing a transformation, with electromyography (EMG) emerging as a key modality in a rapidly expanding ecosystem of commercial and research-grade devices. To translate the ongoing transformation of wearable neural interfaces into robust scientific insights, advances in the field must be supported by precise simulations that faithfully integrate the physiological processes underlying EMG signals and motor control. Over the past decades, electrophysiological simulation software has itself evolved remarkably: early stochastic signal generators [1–4] have given way to biophysically inspired motor neuron (MN) models [5–8], integrated with anatomically and physiologically realistic neuromuscular frameworks [9–12] capable of generating synthetic EMG signals. This evolution reflects not only a deepening understanding of the interplay between neural control, motor unit (MU) physiology, and EMG generation, but also a shift toward physiologically grounded modeling.

Existing EMG simulation tools [1–5, 9–12] have provided important insights into specific aspects of neuromuscular physiology within their respective scopes. However, as research questions increasingly target interactions across neural control, MU organization, muscle mechanics, and signal generation, there is a growing need for modeling frameworks that integrate these components within a unified physiological loop.

To harness this potential, we introduce *MyoGen* (Fig. 1), an open-source neuromuscular and electrophysiology simulation framework that integrates realistic modeling at multiple scales within a single physiological modeling environment. MyoGen combines biophysically grounded MN populations, anatomically realistic spinal circuitry, proprioceptive feedback, and muscle–electrode interactions, enabling coordinated simulation across the full neuromuscular pathway. By unifying spinal-level neural activity, muscle dynamics, and experimentally recorded EMG signals within a modular and extensible framework, MyoGen enables systematic and reproducible investigation of neuromuscular interactions across levels that are not jointly captured by existing relatively single-level simulation approaches. In this article, we describe the architecture of MyoGen and demonstrate its capabilities through simulations that reproduce key statistical features of human motor neurons, enable the generation and decomposition of high-density surface and intramuscular EMG, capture cortical beta-band modulation of descending neural drive and its impact on MU behavior and force generation, and simulate spinal reflexes.

**Fig. 1:**
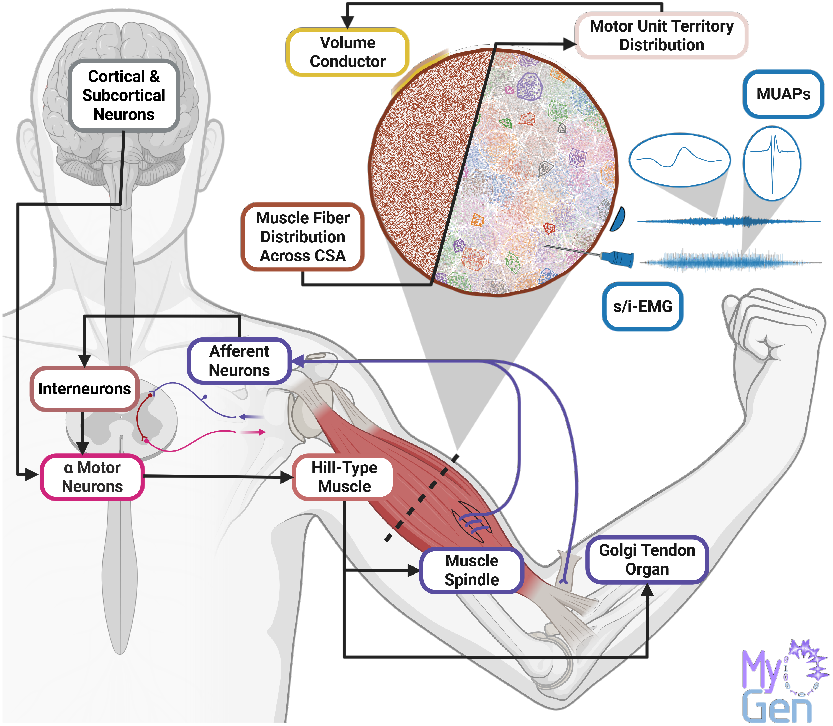
Overview of MyoGen, an open-source framework that simulates the full neuromuscular pathway, from descending neural drive and spinal circuitry to biomechanical muscle models and sensory feedback via muscle spindles and Golgi tendon organs. Motor neurons innervate distributed sets of muscle fibers within the volume conductor, generating motor unit action potentials and synthetic intra-muscular or surface EMG. Together, these components integrate neural, muscular, and electrophysiological processes within a unified modeling framework.

## 2 Results

### 2.1 Physiological framework for MU and EMG simulation

MyoGen reproduces the sensorimotor control loop by integrating descending cortical input, interneuronal circuitry, and afferent feedback at the level of spinal *α*-MN, whose biophysical dynamics generate MU spiking activity, muscle force, and EMG signals (Fig. 1). Proprioceptive feedback from muscle spindles and Golgi tendon organs travels via Ia, II, and Ib pathways, with Ia afferents providing direct input to *α*-MNs and II/Ib afferents interacting with spinal interneurons to modulate motor output (Fig. 1). *α*-MNs can then be driven either by direct current injection, producing mechanistic spike trains that bypass physiological synaptic integration (Fig. 2A), or by synaptic input from a cortical neuron population modeled using independent Poisson or Gamma point processes [13], yielding physiologically realistic MU recruitment and discharge-rate variability (Fig. 2B).

**Fig. 2:**
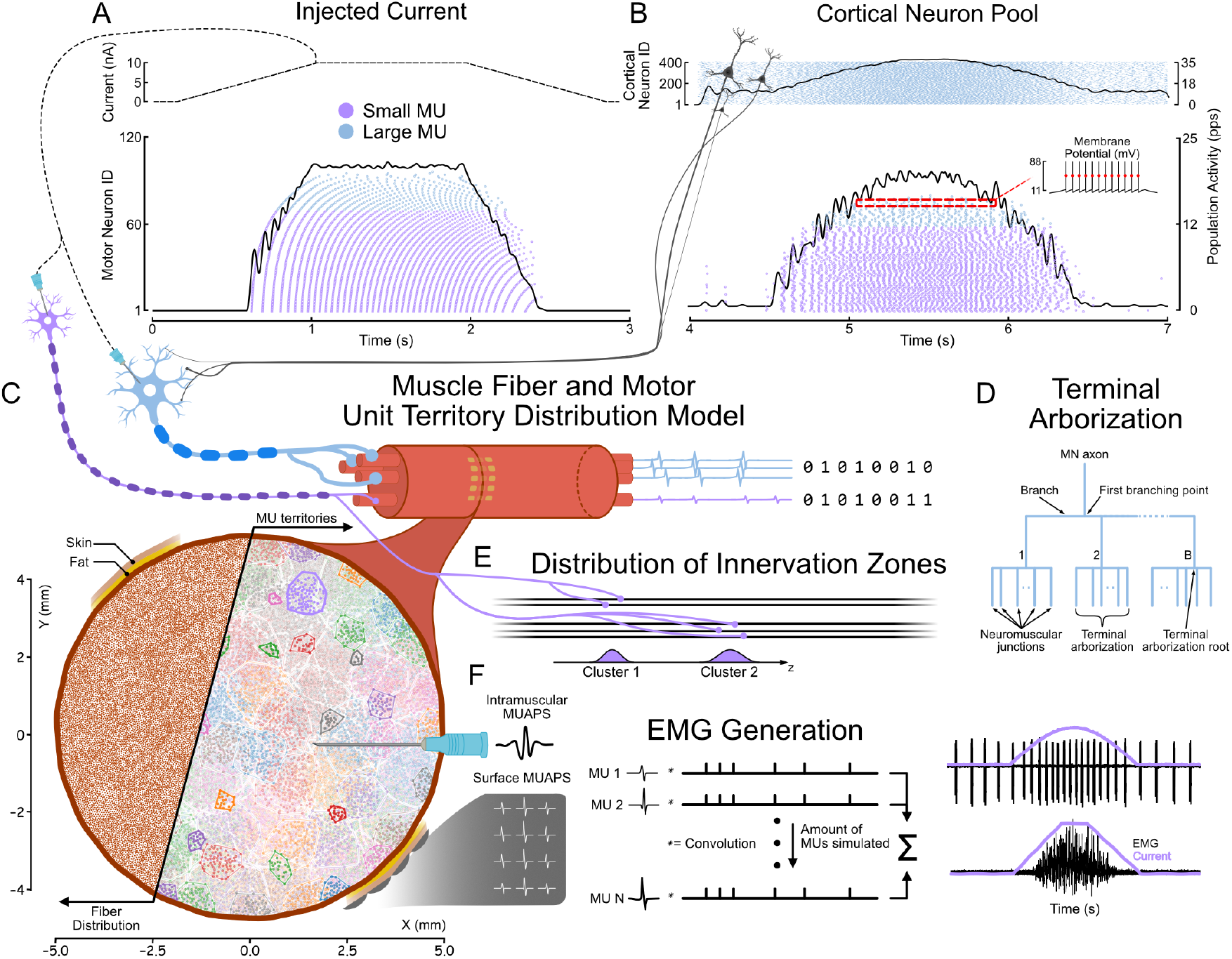
**A. & B**. A pool of *α*-motor neurons (*α*-MNs) can be driven in two ways. In **A**, a direct current is applied to each MN soma in the pool, producing mechanistic discharge patterns that bypass physiological synaptic integration. In contrast, in **B**, the motor neuron pool is driven by a cortical neuron population, resulting in discharge behavior that reflects physiologically structured synaptic input. **C**. The muscle fibers are distributed within the muscle volume and assigned to *α*-MN territories. **D. & E**. The spatial distribution follows a simplified terminal arborization model (**D-E**, reproduced from Konstantin et al. [4]) governing fiber innervation and innervation-zone distribution. **F**. The resulting spike trains are combined with the muscle model to simulate surface and intramuscular EMG signals.

The resulting MU activity drives a physiologically based muscle model that defines the spatial organization of muscle fibers and MUs within the muscle volume (Fig. 2C). MU territories follow a simplified terminal arborization model (Fig. 2D), capturing the experimentally observed [4] longitudinal spread of innervation zones (Fig. 2E). MU action potentials (APs) are then propagated through the surrounding tissue to synthesize intramuscular and surface EMG signals (Fig. 2F). All computational components are implemented in the language or framework most suitable for their function, with biophysical simulations in NEURON [14], performance-critical modules in Cython [15], and higher-level logic in Python. They are connected through standardized data structures based on the Neo framework [16], ensuring compatibility with toolchains such as SpikeInterface [17]. The core API enables each stage of the simulation to be saved and reloaded, avoiding redundant computations when exploring parameter variations. Default parameters are physiologically grounded, allowing users to construct and test complete pipelines without extensive manual tuning. By separating model construction from parameter refinement, the framework maximizes efficiency while maintaining close alignment with experimental workflows. Detailed mathematical descriptions of each simulation step are provided in the Methods.

### 2.2 Decomposable intramuscular and surface EMG simulations

End-to-end simulations demonstrate that the proposed framework generates surface and intramuscular EMG (s/i-EMG) signals that support standard MU decomposition approaches (Fig. 3). These simulations capture the full neural-to-muscle pathway, from descending cortical drive to EMG signal formation.

**Fig. 3:**
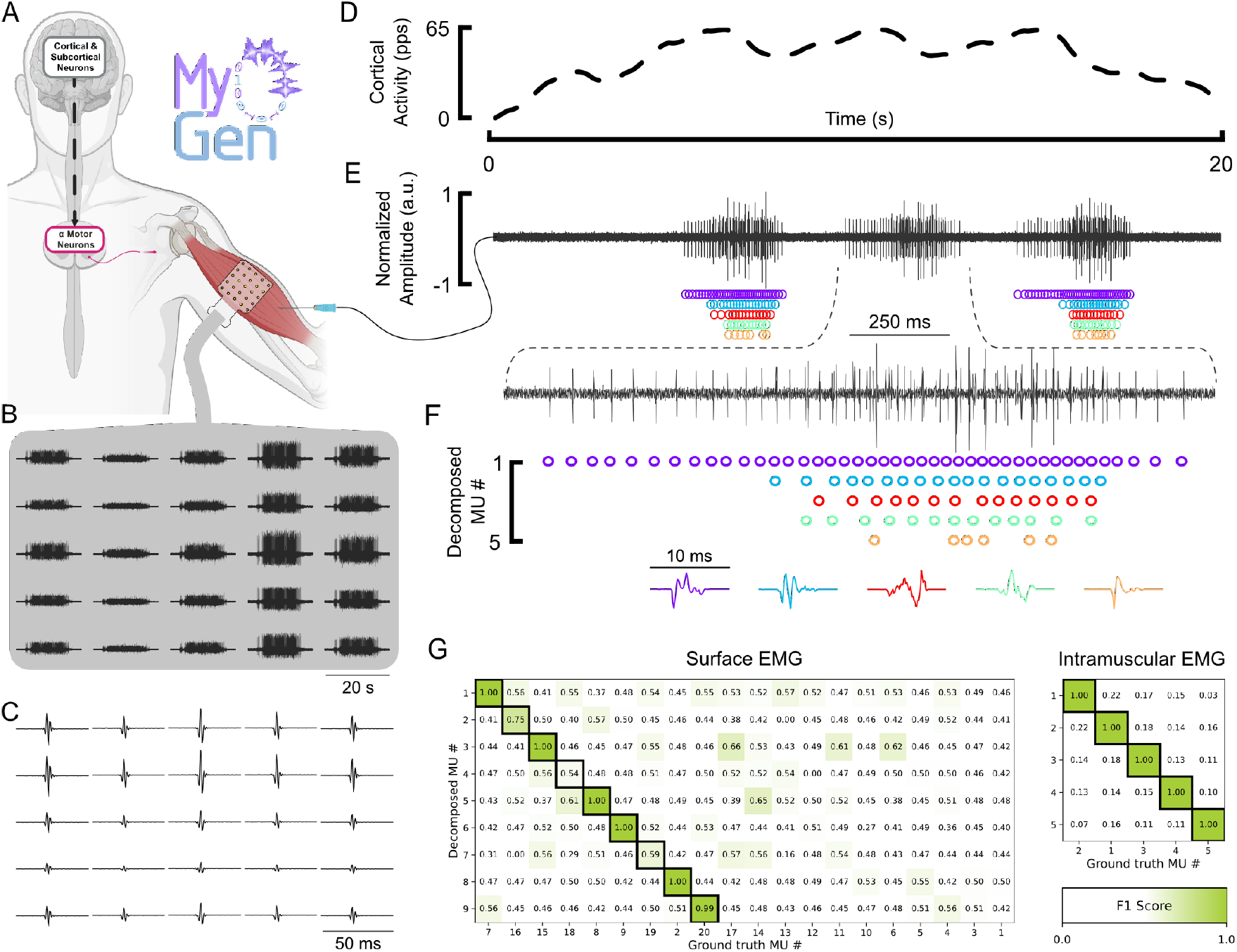
**A**. The minimal EMG simulation pipeline in *MyoGen* comprises three components. Cortical and subcortical neurons projecting to the *α*-MN pool, which can be driven to arbitrary activation levels at specified time points to generate realistic MU discharge patterns (see **D**). The resulting spike trains are then used with a biophysically based muscle volume conductor to synthesize surface or intramuscular EMG (s/i-EMG) signals. **B**. Example of a simulated 5 × 5 electrode grid (2 mm inter-electrode spacing) and the resulting sEMG signals produced by 20 MUs in response to the cortical input shown in **D. C**. Example of a surface MU (MU 1 in **G**) decomposed with MUEdit [18] and subsequently refined with DeMuse [19, 20]. **D**. Cortical drive pattern representing a ramp contraction with a plateau at 65 pps and high variability, generated using Perlin noise. **E**. Example of an iEMG channel comprising five MUs. **F**. Decomposition of the iEMG using EMGLab [21]. **G**. Confusion matrix showing the correspondence between generated and decomposed MUs, quantified using the F1 score (see Methods). The generated (ground truth) MUs are sorted in descending order by their recruitment threshold (1 - the largest MU). Black-bordered cells denote MU matches identified via the Hungarian algorithm.

We simulated a muscle innervated by a pool of 100 MUs and recorded the resulting activity using both a multi-electrode surface grid and a bipolar intramuscular electrode (Fig. 3B–F). The generated EMG signals exhibited physiologically realistic spatial organization, amplitude distributions, and discharge variability, closely resembling patterns observed in experimental recordings.

Surface EMG decomposition recovered a subset of the simulated MU spike trains after refinement, whereas intra-muscular decomposition successfully identified individual MU discharge patterns from a single channel. Decomposition performance was quantified by comparing simulated and reconstructed MU spike trains using the F1 score (Fig. 3G). The resulting confusion matrix revealed clear one-to-one correspondences between simulated and decomposed MUs, with minimal cross-assignment, even in the presence of overlapping MU APs. Together, these results demonstrate that the simulated s/i-EMG signals preserve the structure required for reliable MU decomposition, enabling controlled validation and benchmarking of decomposition pipelines under physiologically realistic conditions.

### 2.3 MU discharge patterns beyond human recordings

To determine whether MyoGen reproduces MU discharge behavior observed in humans, we compared simulated MU populations with experimentally decomposed Mus from three muscles with distinct functional roles: vastus lateralis (VL), vastus medialis (VM), and first dorsal interosseus (FDI). Across muscles, experimentally decomposed motor units defined a characteristic mean discharge rate (DR)–interspike interval (ISI) Coefficient of variation (CV) landscape.

Simulated MU populations generated with MyoGen closely matched this empirical landscape (Fig. 4), with DRs aligned to experimental observations through optimization using Optuna [24]. Across all muscles, simulated MUs occupied the same mean DR–ISI CV space as experimentally decomposed units (Fig. 4B–D), capturing 99.8% of the experimental distribution of mean DRs and 96.8% of the distribution of ISI CVs (Fig. 4B).

**Fig. 4:**
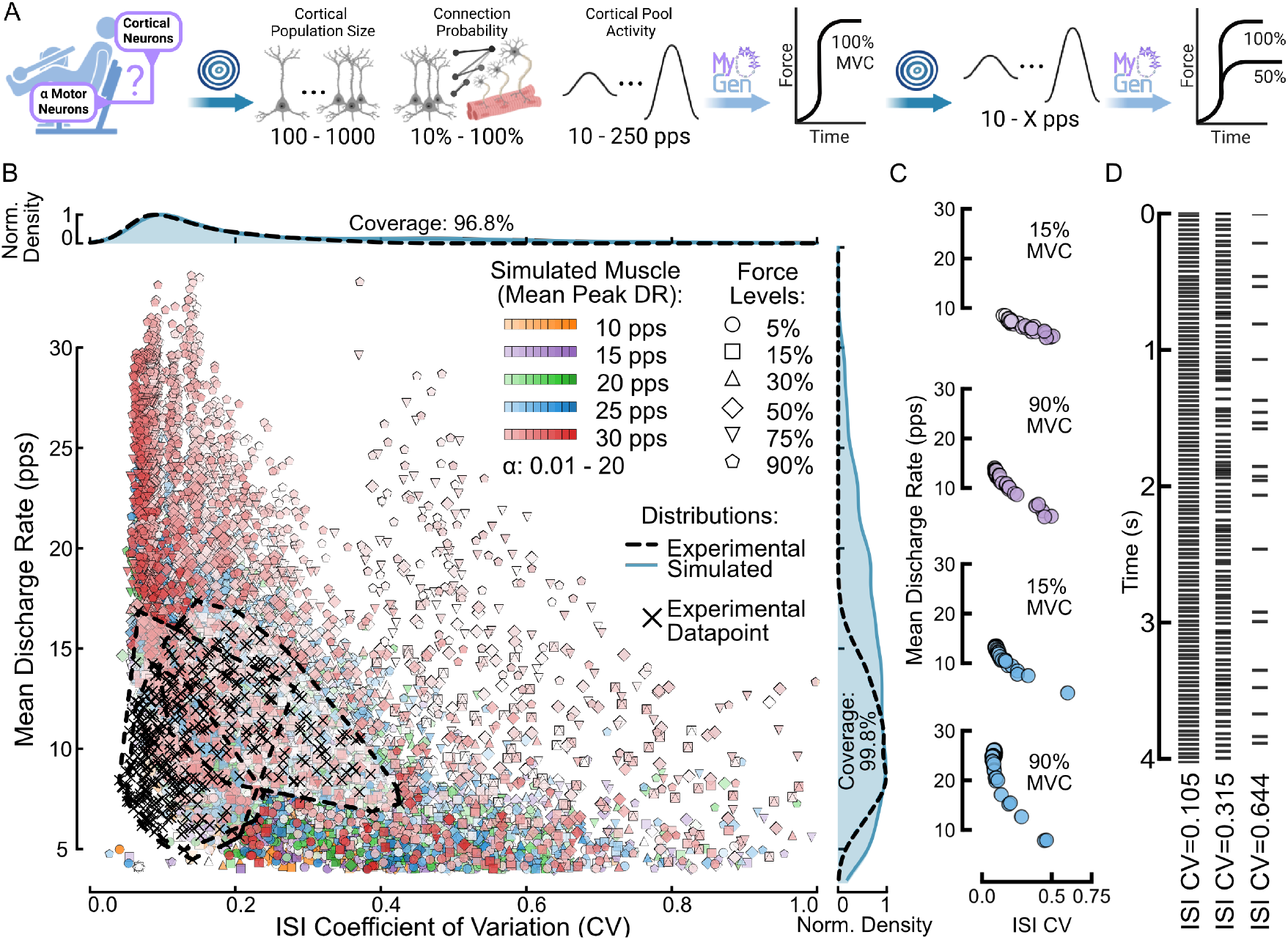
**A**. Most existing motor unit (MU) models rely on feline physiological data [22, 23]. With MyoGen, we simulate MUs with human-like statistics. A pool of 100 *α*-MUs was driven to achieve mean peak discharge rates (DRs) of 10–30 pps by cortical neuron pools modeled as Gamma point processes with fixed shape parameters (see Methods). Using Optuna [24], we optimized cortical pool size (100–1000 neurons), connection probability (10–100%), and overall activity (10–250 pps) to match target FRs. The resulting parameters were used to compute the maximal voluntary contraction (MVC) force [25], followed by a second optimization varying cortical activity to reach 5–90% MVC. **B**. Interspike interval (ISI) Coefficient of variation (CV) versus mean DR for experimentally decomposed MUs (black crosses) and MUs simulated with MyoGen. Normalized marginal distributions are shown above and to the right. Coverage denotes the fraction of the experimental distribution reproduced by the simulations (see Eq. (64) in Methods). **C**. Example MU populations optimized for peak DRs of 15 and 25 pps with a Gamma shape parameter of 1. **D**. Representative spiking behavior of MUs with increasingly stochastic discharge (CV = 0.1–0.6).

Beyond reproducing experimental statistics, the simulations systematically spanned discharge patterns with different degrees of temporal variability, allowing for a controlled exploration of MU recruitment and discharge dynamics that are not accessible in human recordings. They also captured the experimentally observed reduction in CV with increasing force, reproducing human MU behavior across the full range of voluntary muscle contractions.

### 2.4 Recruitment thresholds and resulting electrophysiological parameters

We next examined how recruitment–threshold formulations shape the electrophysiological properties of a MU pool. We generated 100 MUs using either the classical Fuglevand et al. [25] model or the hierarchical formulation of De Luca and Contessa [26], systematically varying the slope parameter from 0.5 to 25. As the slope deviated from 1, recruitment thresholds became correspondingly steeper or shallower (Fig. 5A), inducing structured changes in downstream discharge behavior.

**Fig. 5:**
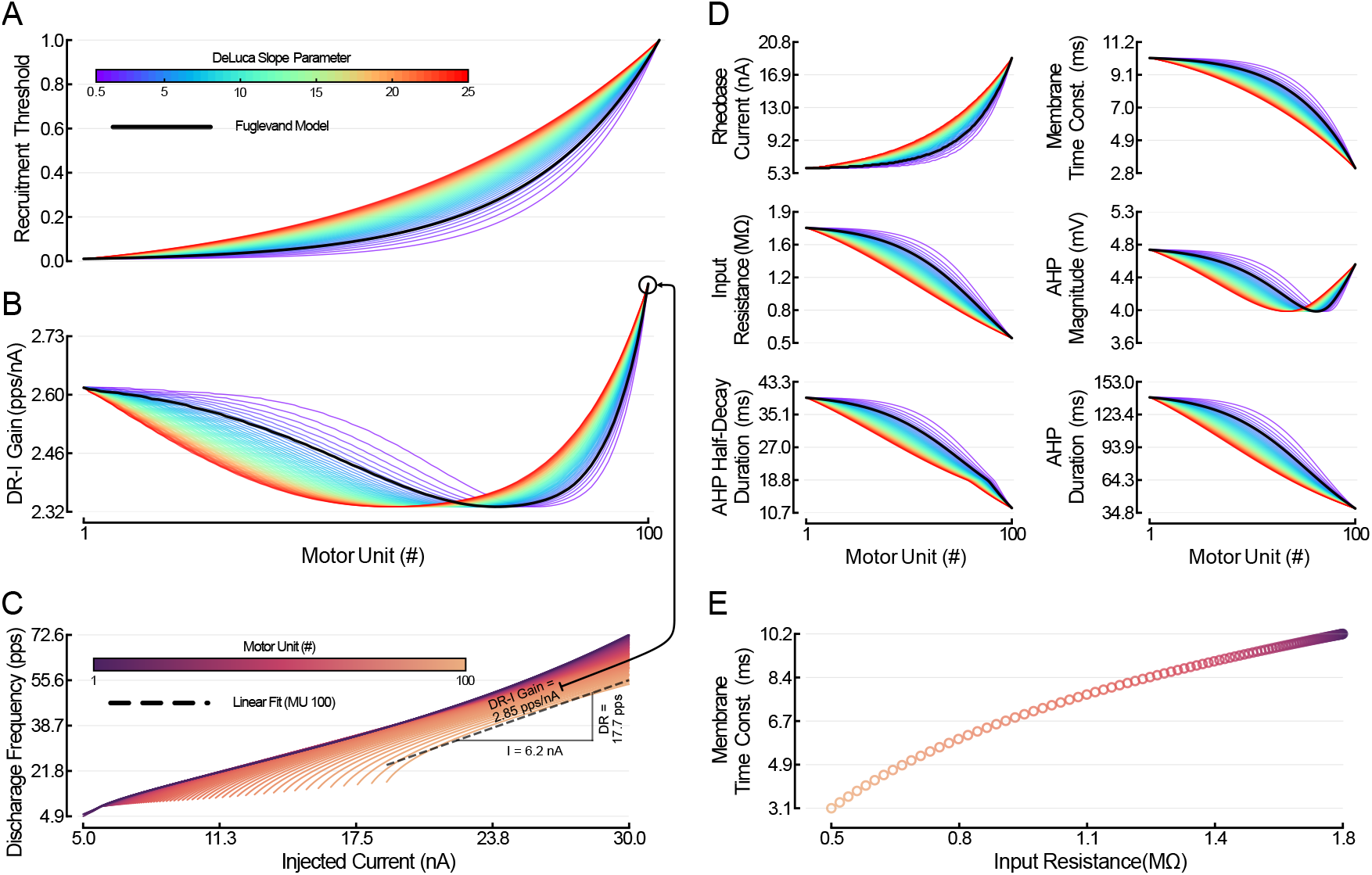
**A**. Recruitment thresholds for 100 motor units (MU) generated using the De Luca and Contessa [26] model with varying slope parameters (color scale), compared with the standard Fuglevand et al. [25] model (black). **B**. Resulting discharge rate - current (DR–I) gains for each motor unit across the pool. **C**. DR–I curves for all motor units, from low- to high-threshold (purple to orange). The linear fit for MU 100 highlights an DR–I gain of approximately 2.85 pps/nA. **D**. Corresponding electrophysiological parameters derived for each MU as a function of recruitment threshold and input–output properties: rheobase current, membrane time constant, input resistance, afterhyperpolarization (AHP) magnitude, AHP half-decay duration, and full AHP duration. **E**. Relation between membrane time constant and input resistance across the MU pool. **D–E**. Across all simulated MUs, parameter values fell within experimentally reported ranges (Table S1), with mean deviations below 3% relative to literature benchmarks.

These threshold manipulations produced systematic shifts in the discharge–intensity (DR–I) gain across the MU pool (Fig. 5B). DR–I gain was estimated by driving each MU with a 5–30 nA current ramp and fitting a linear model to the resulting DR–I relationship (example shown for MU 100 in Fig. 5C), revealing a graded redistribution of excitability across motor units.

From these simulations, we extracted canonical electrophysiological parameters, including rheobase current, input resistance, membrane time constant, and afterhyperpolarization (AHP) magnitude and duration (Fig. 5D). Across all simulated motor units, parameter values fell within experimentally reported ranges, with mean deviations below 3% relative to literature benchmarks (Table S1).

### 2.5 Force modulation under beta-band cortical drive

To assess MyoGen’s ability to reproduce previously reported neuromuscular dynamics, we implemented the cortical oscillation paradigm originally described by Watanabe and Kohn [27] within the MyoGen framework (Fig. 6).

**Fig. 6:**
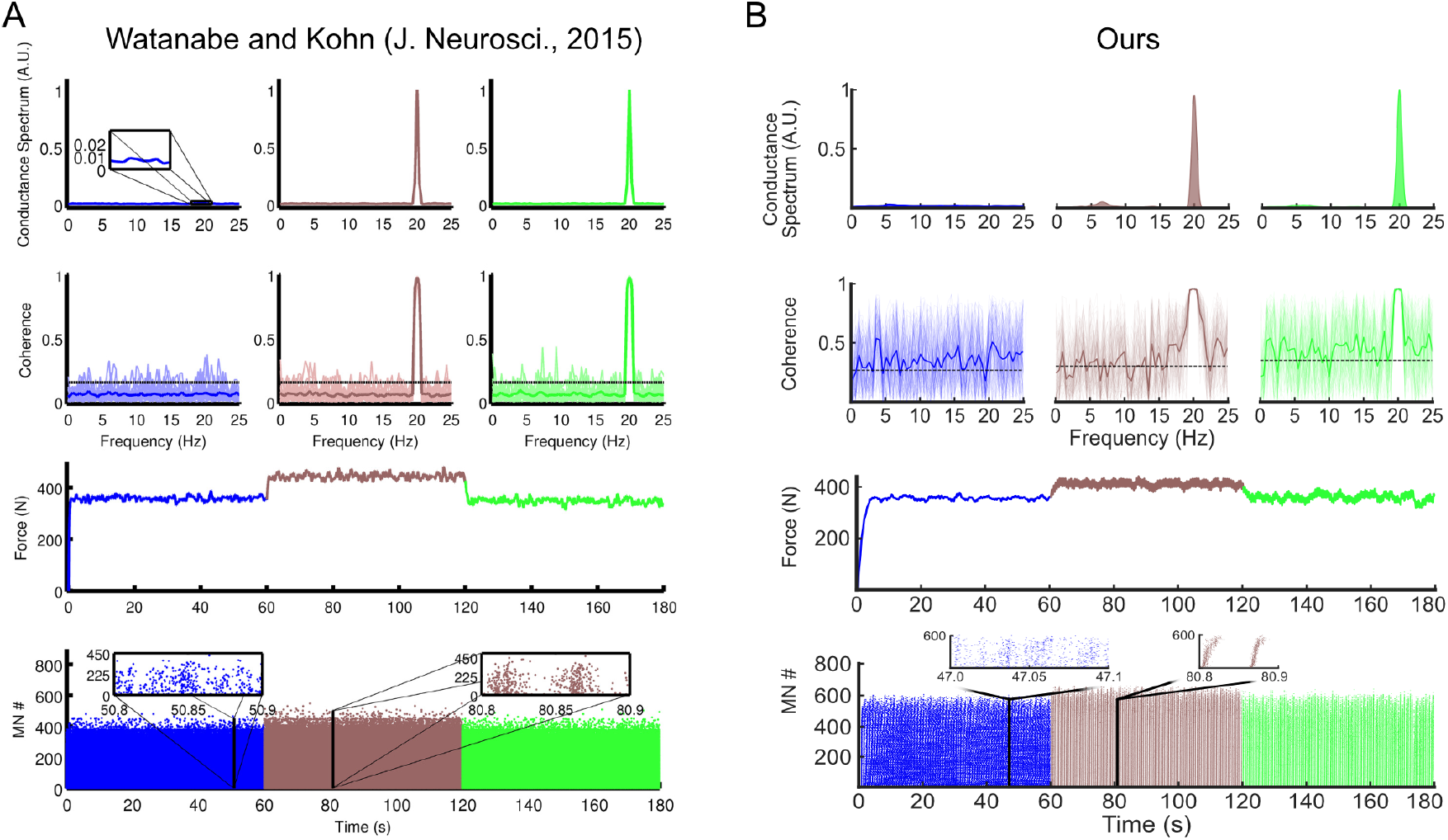
**A**. Cortical oscillation paradigm originally reported in Fig. 3 of Watanabe and Kohn [27] (reproduced with permission). Introducing a 20 Hz oscillatory component to the descending drive increases mean muscle force and induces coherent 20 Hz activity in the output, demonstrating that spinal motoneurons can transform oscillatory cortical inputs into sustained force through nonlinear recruitment mechanisms. From top to bottom: power spectra of the synaptic input, showing the emergence of a 20 Hz peak under oscillatory drive; input–output coherence, revealing frequency-specific coupling; muscle force traces, illustrating an increase in mean force without a change in average discharge rate; and motoneuron rasters, showing intermittent recruitment of higher-threshold units underlying force enhancement. **B**. Implementation of the same cortical oscillation paradigm within MyoGen. Despite differences in model structure and biophysical detail, MyoGen reproduces the key qualitative features of the original neuromuscular regime, including force modulation under oscillatory drive (see force model in Fuglevand et al. [25]), frequency-specific coherence, and recruitment-driven force enhancement. Qualitative differences relative to the original study arise primarily from the use of active dendritic compartments in the motor neuron model (see Methods).

We simulated a pool of 900 MNs innervated by a cortical drive comprising three consecutive phases of 60 s each. In the first phase, the cortical discharge rate was set to 65 pulses per second (pps). In the second phase, a 20 Hz oscillatory component was added to the same baseline discharge rate. In the third phase, the baseline rate was reduced to 58 pps to match the force level of the first phase, following the structure of the original study. Each MN additionally received an independent Poisson noise drive (mean interspike interval, 8 ms).

In contrast to the model of Watanabe and Kohn [27], which employed passive dendrites, MyoGen uses active dendritic compartments throughout, reflecting a more biophysically detailed representation of MN electrophysiology.

The discharge rate of the presynaptic cortical neuron pool was set to 40 pps, consistent with the scaling required to elicit the target MN discharge rates in this model. For the third phase, force output was optimized using Optuna [24] to closely match the force (see force model in Fuglevand et al. [25]) produced during the first phase, analogous to the procedure used in Fig. 4.

The goal of this comparison was not to numerically reproduce the original simulation, but to test whether MyoGen reproduces the same qualitative neuromuscular regime under analogous drive conditions. Similar to the original study, we observed that a beta-band (20 Hz) oscillation at the input of the MN pool increases the average muscle force by recruiting additional *α*-MNs. This demonstrates the non-linear behavior of the motor pool, as the higher-frequency oscillations are mapped into lower-frequency components within the muscle force bandwidth.

### 2.6 Closed-loop spinal circuit simulation

We demonstrate MyoGen’s capability to simulate closed-loop neuromuscular dynamics by implementing a sensorimotor circuit (Fig. 7A) comprising cortical neurons (Fig. 7B), proprioceptive afferents and interneurons (Fig. 7C), *α*-motor neurons (Fig. 7D), a Hill-type musculotendon unit (Fig. 7E), and validated models of the Golgi tendon organ and muscle spindle (Fig. 7F).

**Fig. 7:**
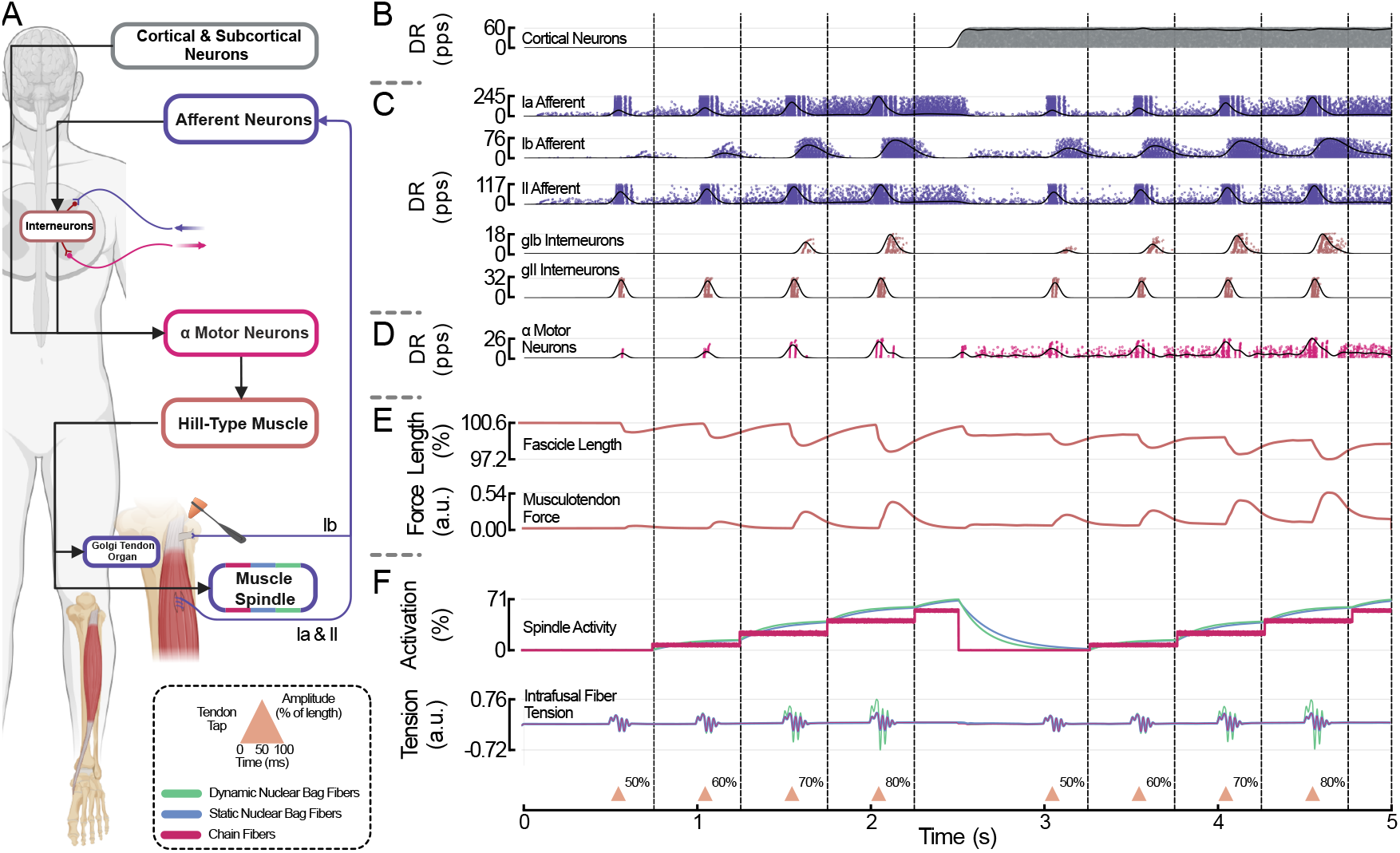
**A**. Schematic of the modeled sensorimotor loop, including cortical and subcortical neurons, afferent pathways (Ia, Ib, II), spinal interneurons, *α* motor neurons, a Hill-type muscle model, the Golgi tendon organ, and the muscle spindle with dynamic bag, static bag, and chain fibers. **B.–D**. Population discharge rates (DR) of cortical neurons (**B**.), Ia, Ib, and II afferents together with interneurons of the Ib and II pathways (gIb and gII) (**C**.), and *α* motor neurons (**D**.). Cortical drive is introduced at *t* = 2.5 s, producing increased motor output and afferent feedback. **E**. Musculotendon mechanics: fascicle length and musculotendon length. Each tendon tap, delivered every 0.5 s at 50, 60, 70, and 80% muscle-length intensity (and restarting after *t* = 2.5 s), produces a rapid mechanical perturbation followed by the corresponding proprioceptive responses. **F**. Muscle spindle responses: overall spindle activation and intrafusal fiber tensions for dynamic bag, static bag, and chain fibers. *γ* drive begins at *t* = 0.75 s and increases in 25 pps increments every 0.5 s (dashed black vertical lines), producing graded modulation of spindle sensitivity. Tendon taps evoke brief high-frequency bursts in Ia and II afferents and oscillations in intrafusal tension.

We simulated the patellar reflex by incorporating three interacting perturbations: periodic tendon taps, graded *γ*-drive modulation, and delayed cortical input.

Tendon taps occurred every 0.5 s and generated brief mechanical perturbations whose amplitudes increased from 50% to 80% of muscle length. Each tap was modeled as a 100 ms triangular ramp, peaking at 50 ms. The graded *γ*-drive, introduced at 0.75 s, increased in 25 pps increments every 0.5 s (Fig. 7F), thereby modulating the sensitivity of the spindle’s static and dynamic components and shaping afferent responses to each tap. When cortical input was introduced at 2.5 s, both the tap amplitude and *γ*-drive were reset to their initial values to isolate the effect of descending drive on the circuit’s dynamics.

### 2.7 Low-dimensional manifolds extraction from simulated agonist–antagonist EMG activity

As a final demonstration of MyoGen’s capabilities, we examined whether systems governed by low-dimensional neural control can be simulated and whether their underlying structure can be recovered from the resulting EMG signals.

We first simulated a simplified paradigm in which the *α*-motor neuron (*α*MN) pools of each muscle group were driven by sinusoidal direct current injections (1 Hz, 20 s recordings; Fig. 8A). Spiking activity closely tracked the injected currents, producing EMG signals across multiple channels (every eighth channel plotted; Fig. 8B). Dimensionality reduction using principal component analysis (PCA) revealed that individual movement cycles occupied distinct trajectories in principal component space, with median trajectories capturing the rotational (cyclical) structure across cycles (Fig. 8C & D). Applying uniform manifold approximation and projection (UMAP) [28] to the same EMG data further confirmed the presence of a low-dimensional mani-fold underlying agonist–antagonist coordination (Fig. 8E & F).

**Fig. 8:**
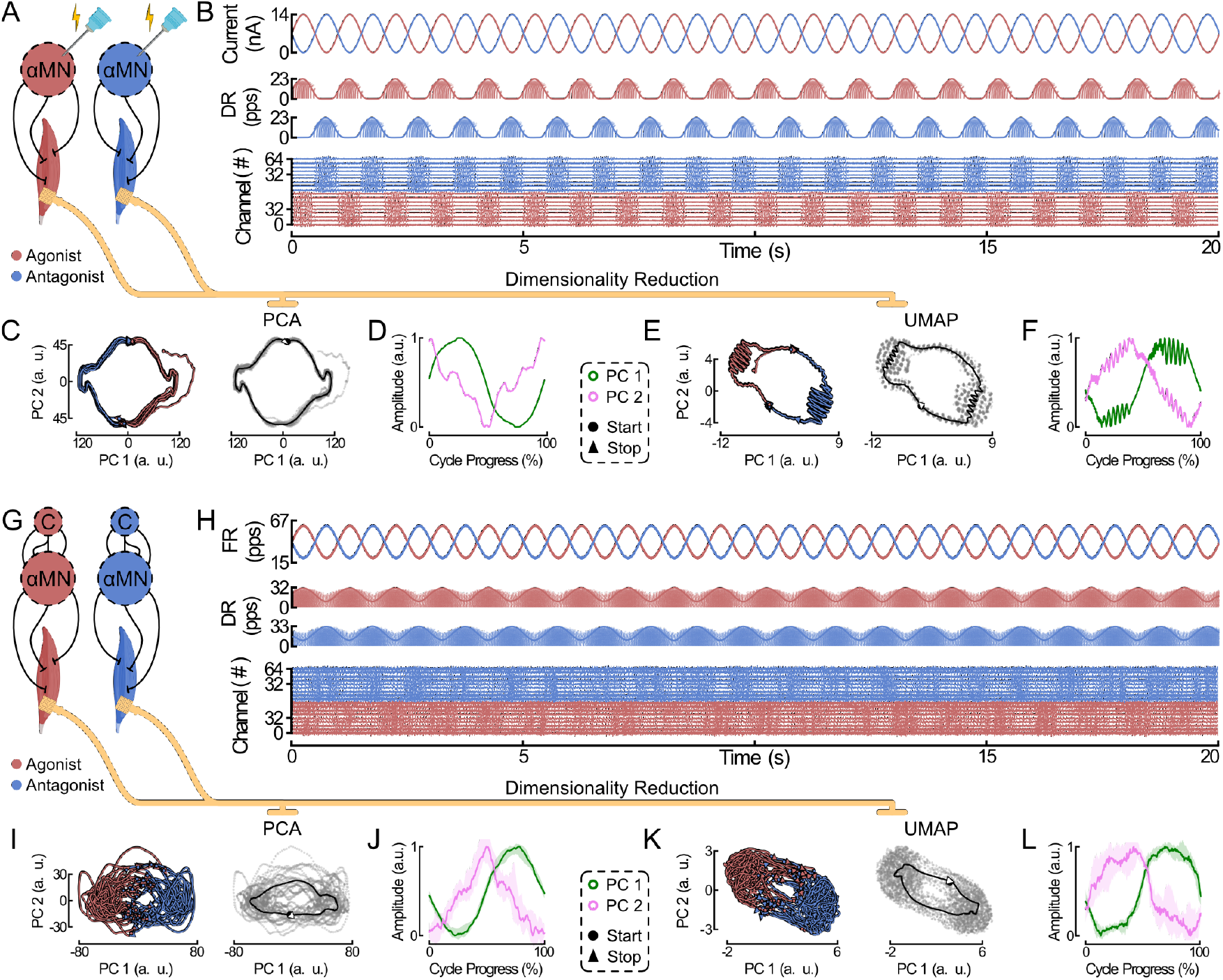
**A–L**. Functional manifold extraction from agonist–antagonist EMG activity under two distinct neural drives. **A**. Agonist and antagonist muscles driven by direct current injection to the *α*-motor neuron (*α*-MN) pools. **B**. Simulation results for **A** shown at successive stages (top to bottom): injected current, spiking activity of the *α*-MNs, and the resulting EMG channels (every eighth channel plotted out of 64 per muscle). **C**. Principal component analysis (PCA) applied to the EMG data from **B**. Left, trajectories from individual cycles, colored by agonist or antagonist activity. Right, median trajectory with all individual data points overlaid in gray. **D**. Principal components (PCs) extracted from the PCA embeddings shown in **C**, averaged over the movement cycle. **E**. Uniform manifold approximation and projection (UMAP) applied to the EMG data from **B. F**. UMAP components corresponding to **E**, averaged over the movement cycle [28]. **G–L**. Same experimental paradigm as in **A**, but with a more physiologically realistic cortical input instead of direct current injection. A cortical neuron pool innervates the *α*-MNs with a connection probability of 30%. Cortical spike trains are modeled as first-order Poisson point processes.

We then extended this paradigm to a more physiologically realistic cortical drive. Cortical neurons were connected to the *α*MN pools with a 30% probability, and spike trains were generated as first-order Poisson point processes (Fig. 8G). Despite increased variability relative to direct current drive, both PCA and UMAP analyses revealed well-defined trajectories and manifolds (Fig. 8H–L).

## 3 Discussion

MyoGen (Fig. 1) builds on decades of influential neuromuscular and electrophysiological modeling [1–5, 9–12] by introducing a framework designed from the outset for modularity and community use. Rather than redefining established approaches, MyoGen extends them through a Neo-based architecture [16] that ensures seamless compatibility with widely adopted analysis ecosystems such as SpikeInterface [17]. This emphasis on interoperability reflects a broader maturation in the field, where long-standing biophysical and computational insights can now be integrated within shared, reusable infrastructures. Through extensive documentation, tutorials, and flexible APIs, MyoGen lowers the barrier to incorporating realistic physiological components into diverse research pipelines, while also supporting full end-to-end simulations when needed. The framework additionally provides pedagogical opportunities: by directly linking mechanistic parameters to observable signals, it offers an intuitive platform for teaching motor control, biophysics, and systems neuroscience. Finally, by enabling controlled generation of ground-truth data across the neuromuscular pathway, MyoGen contributes reproducible benchmarks that complement experimental datasets and support rigorous validation of emerging analytic and machine-learning approaches.

MyoGen emerged from a set of opportunities revealed across the EMG simulation literature [1–5, 9–12]. Foundational frameworks in this space have each advanced a particular aspect of the neuromuscular system: volume-conduction models [1–4] established the principles governing EMG generation; anatomically detailed approaches [9– 11] expanded biophysical realism and tissue specificity; and recent biophysically based platforms [5, 27] have integrated electrophysiological components at high fidelity.

Because these systems were developed to address distinct and often complementary scientific questions, they naturally evolved with different data structures, modeling assumptions, and programming interfaces. While each has proven highly effective within its intended scope, these differences make it difficult to combine existing models into unified, multi-scale workflows or to perform cross-modal validation studies spanning surface and intramuscular EMG, often requiring substantial manual effort and careful reconciliation of assumptions such as parameter definitions and units. In parallel, advances in electrophysiological recording and anatomical characterization have greatly expanded the depth and precision of available datasets, creating a timely opportunity for frameworks that do not replace these established approaches, but instead integrate and coordinate them within a single, coherent computational environment.

MyoGen is designed to meet these emerging needs by providing an open-source architecture that connects spinal circuitry, motor-unit physiology, biophysical volume conduction, and musculotendon dynamics modeling through one extensible API. With MyoGen we allow researchers to reuse existing models while enabling end-to-end, multimodal simulations that support reproducible analysis, benchmarking, and hypothesis testing (Fig. 2, 3, 6, 7, and Fig. 8).

To support this integrative and reusable design, comprehensive documentation is essential for reproducible neuromuscular research. We provide extensive API documentation with type-safe interfaces, worked examples spanning basic to advanced use cases, and guides to the Neo Block data structures. Together, these resources enable researchers to reliably reproduce published results and extend the toolkit to novel applications. In line with FAIR principles [29] for research software, MyoGen incorporates machine-readable type annotations, API documentation with cross-referencing, and structured example galleries, making the toolkit Findable, Accessible, Interoperable, and Reusable.

The integrated simulation of the complete neuromuscular pathway enables investigations that naturally extend from the complementary contributions of prior models. The ability to simulate both surface and intramuscular EMG (Fig. 2) allows direct comparison of signals derived from the same motor-unit populations, offering a controlled platform for optimizing recording strategies and interpreting experimental measurements. In addition, MyoGen produces MU populations with human-like discharge statistics (Fig. 4 & Fig. 5), allowing simulations to capture the variability observed in experimental datasets (Table S1). By incorporating a comprehensive spinal network (Fig. 7), the framework extends beyond feedforward activation to encompass closed-loop motor control, creating opportunities to investigate how cortical neural manifolds are transformed by spinal circuitry into force or kinematics manifolds.

In Fig. 8, we demonstrate that low-dimensional structure can be extracted from simulated surface EMG during an agonist–antagonist task. The resulting manifolds show low-dimensional structure and task-dependent trajectories consistent with those we previously reported from experimental recordings in motor-complete spinal cord–injured [30] and uninjured participants [31, 32]. These results suggest that MyoGen provides a suitable platform for investigating how neuromuscular dynamics give rise to structured EMG activity under both healthy and pathological conditions.

Beyond this, MyoGen offers a flexible framework for future studies aimed at dissecting the neural origins of low-dimensional motor representations. In particular, the simulator enables systematic investigation of whether muscle-level synergies or motor-unit–level synergies provide a more appropriate description of cortical control, a question that remains actively debated in the literature [33]. While recent evidence supports the existence of motor-unit–level synergies [34], their functional role and dependence on neural drive, recruitment dynamics, and task constraints are not yet fully understood. These questions open avenues for future investigations that build on the MyoGen framework to further elucidate the neural organization of motor control. A central challenge in biophysical neuromuscular modeling is the principled identification of parameter sets that reproduce experimentally observed neural and muscular dynamics. Previous studies have achieved realistic motor neuron and muscle behavior by exploring vast parameter spaces through manual adjustment, trial-and-error, or high-performance computing, sometimes running millions of simulations across supercomputers [35, 36]. MyoGen provides a complementary strategy by integrating Bayesian optimization via Optuna [24], enabling automated, data-driven calibration of high-dimensional models against multiple physiological observables. This approach allows simultaneous tuning of descending drive parameters to match motor neuron discharge statistics, discharge variability, and force output, all without manual intervention or extensive computational infrastructure. As a result, MyoGen facilitates the construction of individualized, reproducible models, supports systematic comparisons across muscles and contraction levels, and enables scalable studies of inter-individual variability and pathological motor control, opening new possibilities for both research and clinical applications.

Despite these advances, several considerations warrant attention. The current muscle geometry implementation relies on simplified cylindrical representations, which may not fully capture the anatomical diversity of specific muscles. Future versions could incorporate more detailed geometries derived from medical imaging data [9], further enhancing biophysical realism. Large-scale simulations involving thousands of motor units remain computationally demanding, which may limit accessibility for users without high-performance computing resources; ongoing optimization efforts aim to improve efficiency and enable cloud-based deployment. While MyoGen is modular and extensively documented, fully exploiting its capabilities may require familiarity with Python, computational modeling, or neurophysiology. Ongoing development of user-friendly interfaces and interactive tutorials aims to further democratize access, making the framework accessible to a broader community of researchers and educators. Finally, the present implementation primarily represents healthy MU populations, with pathological conditions currently modeled through parameter adjustments rather than explicit disease-specific mechanisms. These avenues highlight opportunities for continued development and expansion, ensuring that MyoGen remains a versatile and broadly applicable platform for both fundamental and translational investigations.

## 4 Conclusion

Neuromuscular research is constrained by the complexity of integrating neural control, motor unit physiology, muscle mechanics, and electrophysiology. MyoGen provides an open-source platform that unifies biophysically informed spinal circuitry, proprioceptive feedback, musculotendon dynamics, and multi-modal EMG generation within a single framework.

The platform reproduces key experimental benchmarks with high fidelity: simulated motor unit populations capture 99.8% of discharge-rate distributions and 96.8% of discharge variability across multiple human muscles, with all electrophysiological parameters within 3% of literature values. Generated surface and intramuscular EMG signals preserve the structure required for decomposition, enabling controlled validation of analysis pipelines under physiologically realistic conditions.

Data-driven calibration via Bayesian optimization with Optuna allows efficient exploration of complex parameter spaces, while a Neo-based architecture facilitates integration with other established analysis pipelines.

By combining reproducibility, modularity, and accessibility, MyoGen complements existing experimental and computational approaches, providing a platform to investigate open questions in motor control, including the neural basis of muscle synergies, cortical contributions to force modulation, and mechanisms underlying pathological motor output.

## 5 Methods

### 2.1 MyoGen framework architecture

The MyoGen framework (Fig. 1 and Fig. 9A) was developed to simulate the full neuromuscular control loop, spanning descending cortical drive, afferent feedback, and the mechanisms underlying muscle force and EMG generation. Its modular architecture allows users to independently configure, execute, and interconnect physiological components while preserving computational efficiency and reproducibility.

**Fig. 9:**
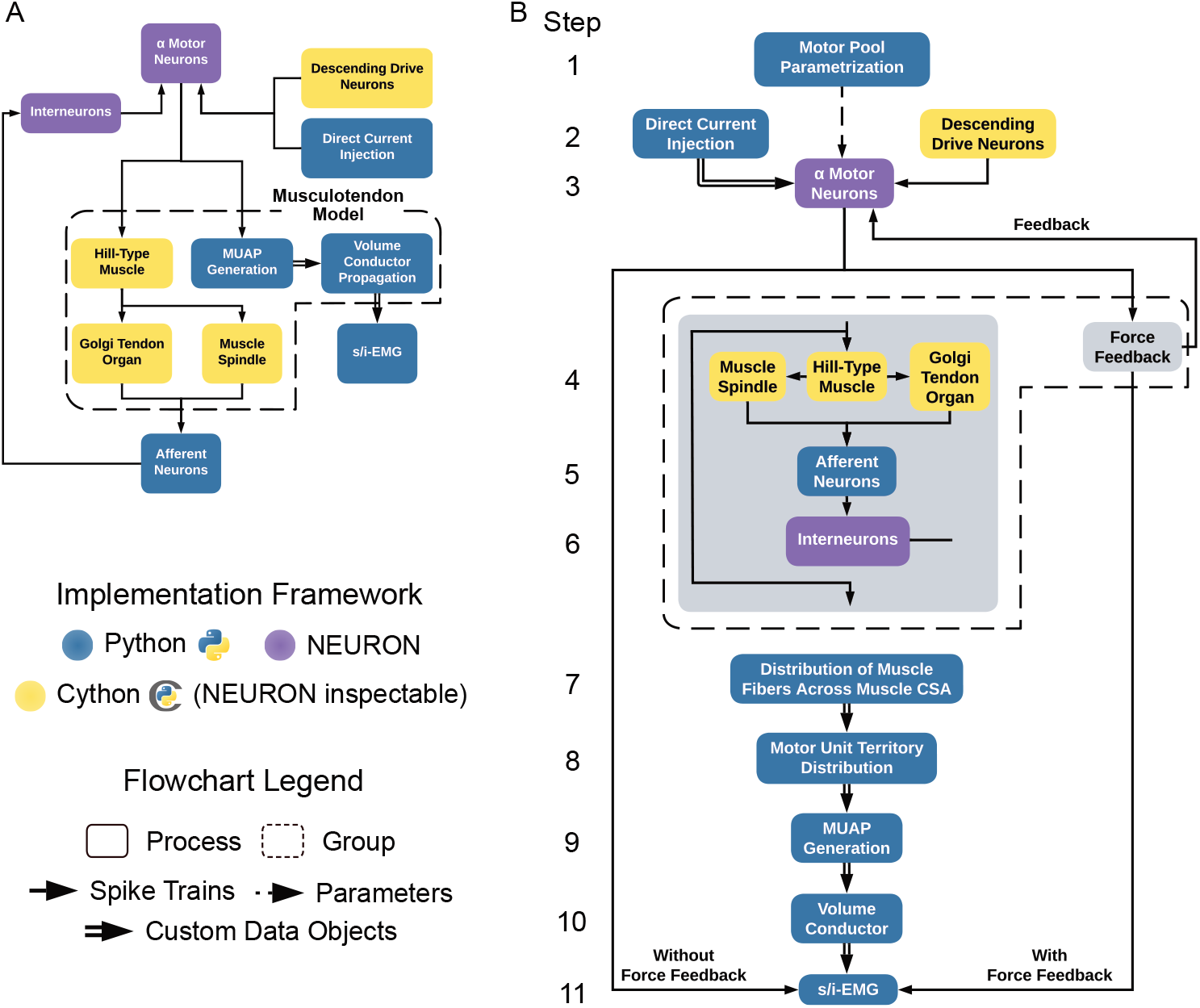
**A**. Flowchart of the *MyoGen* framework showing all available components and their interactions. The output of each component can be saved and reloaded to bypass unnecessary recalculations when adjusting only part of the simulation. Components are color-coded by implementation framework: blue for *Python*, yellow for *Cython*, and purple for *NEURON*. This architecture enables customization of closed-loop simulations, such as coupling the Hill-type muscle, spindle, and Golgi tendon organ models to provide feedback to spinal neurons. **B**. Timeline overview of the EMG simulation pipeline with and without force feedback. Horizontally aligned blocks are computed in parallel.

Communication between components is mediated through standardized data objects representing spike trains, parameters, and system states. Simulation outputs can be serialized and reloaded, enabling redundant computations to be skipped when only subsets of the model are modified. This design enables rapid prototyping and closed-loop experimentation without requiring recomputation of the entire simulation cascade.

To balance biophysical realism with computational scalability, MyoGen combines three complementary implementation frameworks:

- **Python 3.12** (blue modules) provides high-level orchestration, parameter management, and data handling, coordinating the simulation pipeline and managing persistent storage of intermediate results.
- **Cython 3.1.4** [15] (yellow modules) is used for performance-critical components, including the Hill-type muscle model, muscle spindle, and Golgi tendon organ (GTO) feedback. These modules remain observable within NEURON, ensuring consistent state access across frameworks.
- **NEURON 8.2.6** [14] (purple modules) implements biophysically detailed models of *α*-motor neurons.

### 5.2 Cortical neurons representing the descending drive

The descriptive statistics and data on *α*-MN presynaptic processes related to premotoneuronal pathways and voluntary drive remain scarce, as does the characterization of the diversity and complexity of these inputs. To address this, descending drive spike trains are modeled as non-homogeneous Gamma renewal point processes [37], wherein each spike occurrence depends solely on the preceding event, rendering interspike intervals (ISIs) statistically independent and identically distributed.

These presynaptic processes were implemented as an integrate-and-fire point neuron model where the instantaneous drive *y*(*t*) accumulates in a state variable *y*_*i*_(*t*) over discrete time steps Δ*t* (Eq. 1). Spike emission occurs when this cumulative variable reaches a stochastic threshold *θ* (Eq. 2), sampled from the product of *N* independent uniform random variables *U*_*i*_~𝒰 (0, 1) (Eq. 3). This construction exploits a fundamental property of probability theory: the sum of *N* independent exponential random variables yields an Erlang-distributed variable of order *N*, thereby conferring precise control over ISI variability through the shape parameter *N*. Following each event, *y*_*i*_(*t*) resets to zero and a new threshold is drawn, preserving interval independence across successive events.

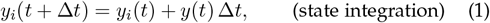

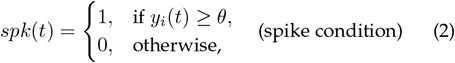

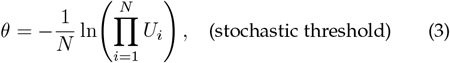

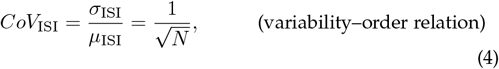

The parameter *N* governs the temporal regularity of spike trains: at *N* = 1, the model reduces to a Poisson process with unit coefficient of variation (*CoV*_ISI_ = 1), whereas for *N >* 1, variability decreases according to 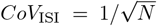 (Eq. 4). This behavior reflects temporal summation dynamics analogous to a neuron integrating *N* stochastic inputs arriving at exponentially distributed intervals. Whether the process is homogeneous or inhomogeneous is determined exclusively by the modulatory signal (*y*(*t*)). A constant *y*(*t*) yields a time-invariant rate and thus a homogeneous process, whereas any temporal variation in *y*(*t*) imposes a time-dependent rate and produces an inhomogeneous process. While the superposition of independent non-Poisson processes does not strictly generate Poisson statistics [38], such frameworks provide tractable and biologically plausible representations for evaluating neuronal responses to random synaptic bombardment.

### 5.3 Biophysical neuron modeling

The neuron models employed in this study (Fig. 4 & 5) describe the dynamics of the electrical potentials along the membrane of excitable cells. The governing equations are based on core conductor and cable theory [39]. Each compartment is morphologically represented by a cylinder and an equivalent electrical circuit comprising passive elements (electrotonic properties) and active elements (voltage-dependent ion channels). The specification of morphological and electrotonic parameters, together with the ion channel dynamics, aims to capture the most relevant electrophysiological characteristics observed in both animal models and humans.

#### 5.3.1 Morphological and electrotonic properties

The electrical behavior of each neuronal compartment can be described by the discrete-space cable equation, which expresses how the transmembrane potential evolves under the influence of membrane currents and axial coupling. For compartment *j*, the transmembrane potential *v*_*j*_ evolves according to Eq. 5, governed by the membrane capacitance *c*_*j*_, membrane resistance *r*_*j*_, and multiple current sources: synaptic inputs *i*_syn,*j*_, ionic currents *i*_ion,*j*_, and externally injected currents *i*_inj,*j*_. The final term in Eq. 5 accounts for axial currents from adjacent compartments *k*, where *r*_*jk*_ represents the cytoplasmic resistance between compartments *j* and *k*.

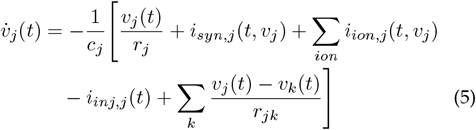

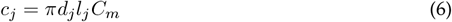

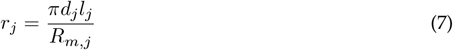

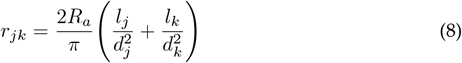

In the present model, the *α*-MN morphology is reduced to two compartments (somatic and dendritic) coupled by a single axial resistance. This simplification removes spatially distributed dendritic propagation, so the cable equation is represented by only two voltage equations, one per compartment. However, the full system remains high-dimensional because each voltage-gated ionic conductance introduces additional coupled differential equations through its activation and inactivation variables. In this reduced morphology, the axial term represents only the bidirectional interaction between the soma and the dendrite.

These compartmental parameters are determined by both the underlying biophysical properties and the geometry of the compartment. The membrane capacitance *c*_*j*_ (Eq. 6) scales with the compartment surface area, being a function of its diameter (*d*_*j*_), length (*l*_*j*_), and the specific membrane capacitance (*C*_*m*_). Similarly, the membrane resistance *r*_*j*_ (Eq. 7) depends on the compartment morphology and its specific membrane resistance (*R*_*m,j*_). The axial resistance *r*_*jk*_ connecting adjacent compartments is defined in Eq. 8 and reflects the specific axial resistance (*R*_*a*_) modulated by the geometry of the connected compartments.

#### 5.3.2 Voltage-gated channels

Beyond these passive properties, the model incorporates active membrane mechanisms that govern the generation of APs and regulate neuronal excitability. These active properties are represented by voltage- and time-dependent ion channel dynamics, where the specific conductance of each ion channel (*g*_*ion*_) evolves according to the local trans-membrane potential *v*_*j*_ and time (*t*). Each ion channel conductance generates a transmembrane current (*I*_*ion*_) whose magnitude and direction are determined by the driving force between *v*_*j*_ and the ion’s equilibrium potential (*E*_*ion*_), as expressed in Eq. 9.

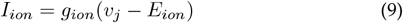

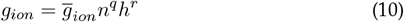

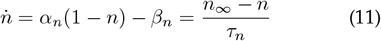

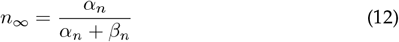

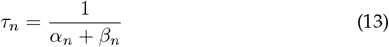

The conductance dynamics are defined through kinetic descriptions involving one or more gating variables, each representing a distinct activation or inactivation process. As shown in Eq. 10, each conductance is scaled by its maximal value 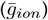, with exponents *q* and *r* indicating the number of independent activation and inactivation gates, respectively. These gating variables evolve according to Hodgkin-Huxley kinetics (Eqs. 11–13), where the generic activation and inactivation variables *n* and *h* (both constrained to [0, 1]) transition between states at voltage-dependent rates *α*_*n*_ and *β*_*n*_ [40].

#### 5.3.3 Synapses

Synaptic transmission is captured through postsynaptic currents (*i*_syn_) determined by the synaptic conductance (*g*_syn_) and the electrochemical driving force between the postsynaptic potential *v*_*j*_ and the synaptic reversal potential *E*_syn_ (Eq. 14). The conductance exhibits bi-exponential kinetics characterized by two state variables, *A* and *B*, decaying with distinct time constants *τ*_1_ and *τ*_2_ (Eqs. 16– 17); the rapid component *A* governs rise dynamics, while the slower component *B* (*τ*_2_ *> τ*_1_) controls decay kinetics (Eq. 15). Each presynaptic spike instantaneously increments both state variables (Eqs. 18–19), with normalization factors *f*_t_ and *f*_g_ (Eqs. 20–21) ensuring proportional scaling of peak conductance amplitude. This formulation enables the computationally efficient integration of multiple convergent inputs—each with its own independent weight and timing—within a unified framework, where 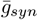 denotes the nominal maximal conductance.

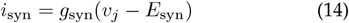

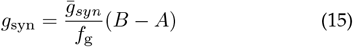

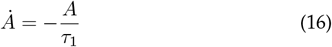

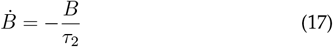

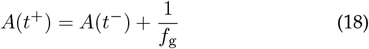

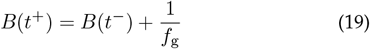

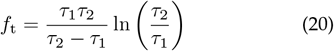

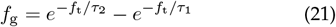

#### 5.3.4 α-Motor neurons

Spinal *α*-MNs were modeled to reproduce essential electrophysiological properties spanning the motor pool, from slow fatigue-resistant (smallest) to fast-fatigable (largest) motor units in the lumbar region of anesthetized cats. Parameterization targets included passive membrane characteristics such as input resistance (*R*_in_) [41] and membrane time constant (*τ*_*m*_) [42], together with active discharge properties including rheobase current (*I*_rheo_), AP amplitude, afterhyperpolarization dynamics [41], frequency–current relations [43], and persistent inward currents (PICs) [7, 43, 44].

MyoGen allows straightforward configuration of motor nucleus properties, enabling flexible control of motor pool size and composition across simulations. We approximated neuronal morphology using reduced bi-compartmental architectures comprising soma and dendrite. We scaled morphological parameters (diameter *d*_*j*_, length *l*_*j*_, and specific membrane resistance *R*_*m*_) across the motor pool to enforce the size principle, as described in Sec. 5.3.5, while keeping passive electrical properties uniform (Table 1). To reproduce force modulation driven by cortical beta-band oscillations, we approximated the soleus musculature using a pool of 900 *α*-motor neurons. To demonstrate physiologically realistic human-like motor unit discharge patterns, we used a reduced pool of 100 *α*-motor neurons. For closed-loop spinal-level simulations and manifold analysis, we adopted a motor pool configuration matching the physiological characteristics of the human FDI muscle [45]. Boundary values for the smallest (*α*-MN1) and largest (*α*-MN120) *α*-MNs are detailed in Table 2.

**TABLE 1:** Specific membrane properties, ionic equilibrium potentials, and excitatory (glutamatergic) and inhibitory (glycinergic) synaptic parameters adopted for the *α*-MN pool [46, 47].

| Symbol | Description | Unit | Value |
| --- | --- | --- | --- |
| $C_m$ | Specific membrane capacitance | $\mu\text{F}/\text{cm}^2$ | 1 |
| $R_a$ | Specific axial resistivity | $\Omega \text{ cm}$ | 70 |
| $E_l$ | Resting potential | mV | 0 |
| $E_{Na}$ | Sodium equilibrium potential | mV | 120 |
| $E_K$ | Potassium equilibrium potential | mV | -10 |
| $E_{Ca}$ | Calcium equilibrium potential | mV | 140 |
| $E_{Glu}$ | Excitatory synaptic reversal potential | mV | 70 |
| $\tau_{1,Glu}$ | Excitatory synaptic time constant 1 | ms | 0.20 |
| $\tau_{2,Glu}$ | Excitatory synaptic time constant 2 | ms | 0.30 |
| $\bar{g}_{Glu}$ | Excitatory max. conductance | $\mu\text{S}$ | 0.050 |
| $E_{Gly}$ | Inhibitory synaptic reversal potential | mV | -10.7 |
| $\tau_{1,Gly}$ | Inhibitory synaptic time constant 1 | ms | 0.25 |
| $\tau_{2,Gly}$ | Inhibitory synaptic time constant 2 | ms | 0.82 |
| $\bar{g}_{Gly}$ | Inhibitory max. conductance | $\mu\text{S}$ | 0.009 |

**TABLE 2:**
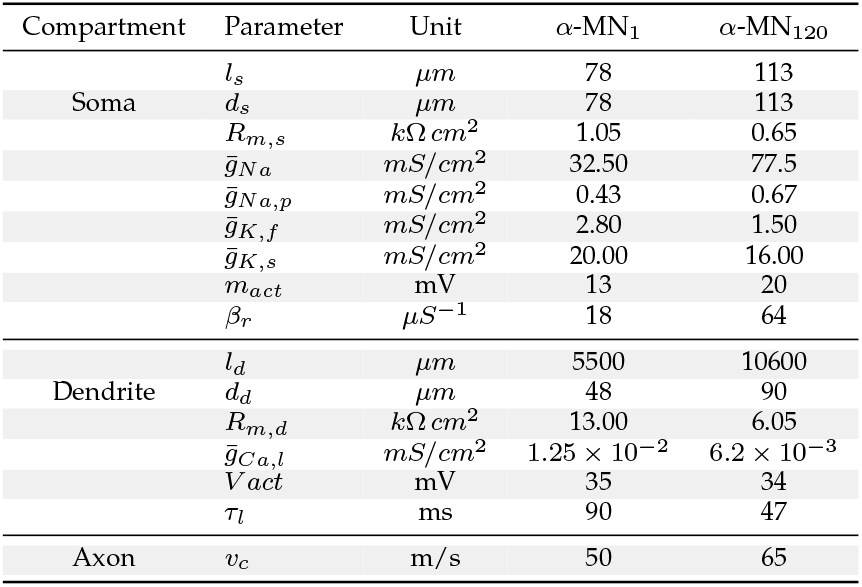
Morphological and electrotonic parameters of the MN population [47].

Active membrane properties were implemented through compartment-specific voltage-gated conductances. The dendritic compartment encompassed an L-type calcium channel (*g*_Ca,l_, Eq. 26) that generates a dendritic PIC, from which synaptic amplification emerges as a physiological consequence. The somatic compartment included the ionic mechanisms underlying AP generation: fast and persistent sodium channels (*g*_*Na*_, *g*_*Na,p*_; Eqs. 22, 23) governing initiation and subthreshold depolarization, complemented by fast and slow potassium channels (*g*_*K,f*_, *g*_*K,s*_; Eqs. 24, 25) controlling repolarization and afterhyperpolarization. Maximal conductances and gating kinetics (Eqs. 27–32) parameters are specified in Table 2, with ionic equilibrium potentials referenced to a normalized resting level of 0 mV (conventional ≈–60 mV shifted to emphasize deviations from baseline) provided in Table 1.

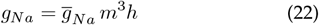

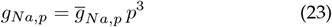

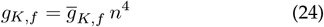

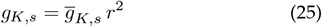

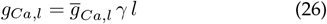

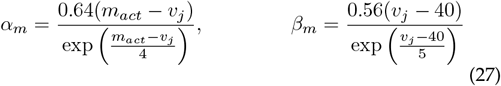

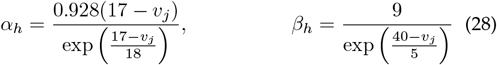

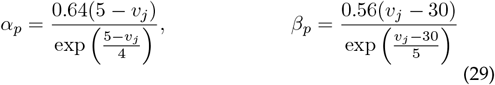

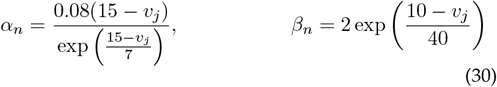

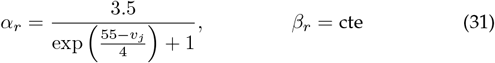

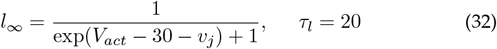

We define the conduction distance as the anatomical path length traveled by APs along motor axons, from the *α*-motor neuron soma in the spinal cord to the neuromuscular junction. As most simulations are not tied to a specific muscle, we treat this distance as a default, configurable parameter representing an average path length to distal limb muscles. We model axonal properties to capture the associated propagation delay. We assume a fixed conduction distance of 0.6 m for all *α*-motor neurons, consistent with the approximate separation between the FDI muscle and its motor nucleus in the cervical spinal cord; only simulations addressing FDI-specific reflex circuitry require the muscle-specific configurations. We scale the conduction velocity (*v*_*c*_) with motor unit size to reflect the well-established relationship between motor axon diameter and conduction velocity (Table 2) [5].

Excitatory synaptic inputs were modeled through glutamatergic conductances (*g*_Glu_ and *E*_Glu_) in the dendritic compartment, representing convergent drive from descending pathways, Ia afferents, and group II interneurons (INs). Inhibitory synaptic inputs, representing group Ib INs, were modeled as glycinergic conductances (*g*_Gly_ and *E*_Gly_) in the somatic compartment [48]. Synaptic kinetics and peak amplitudes (Table 1) were calibrated to reproduce somatic excitatory postsynaptic potentials (EPSPs) [49–52] and inhibitory postsynaptic potential (IPSPs) characteristics observed experimentally in cats [53]. The resulting equivalent electrical circuit of the *α*-MN can be found in Fig. 10.

**Fig. 10:**
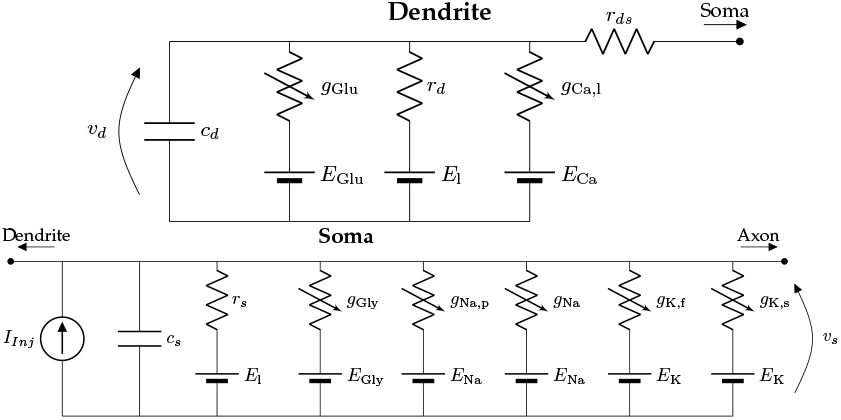
The model comprises dendritic and somatic compartments coupled by an axial resistance (*r*_*ds*_) that governs intracellular current flow between compartments. The dendritic compartment includes a membrane capacitance (*c*_*d*_), a passive leak conductance (*r*_*d*_, *E*_l_), an excitatory synaptic conductance (*g*_Glu_, *E*_Glu_), and an L-type calcium channel (*g*_Ca,l_, *E*_Ca_), which generates the dendritic component of the persistent inward current (PIC). The somatic compartment contains a membrane capacitance (*c*_*s*_), a passive leak conductance (*r*_*s*_, *E*_l_), an inhibitory synaptic conductance (*g*_Gly_, *E*_Gly_), and voltage-gated channels mediating AP depolarization and repolarization: fast and persistent sodium (*g*_Na_, *g*_Na,p_, *E*_Na_) and fast and slow potassium (*g*_K,f_, *g*_K,s_, *E*_K_) conductances. The *g*_Na,p_ contributes to the somatic component of the PIC, facilitates spike initiation, and helps maintain depolarization during the repolarization phase. The somatic voltage (*v*_*s*_) and dendritic voltage (*v*_*d*_) represent local transmembrane potentials, while an injected current source (*I*_Inj_) applies depolarizing or hyperpolarizing input to the soma.

#### 5.3.5 α-Motor neuron pool

Building upon the individual *α*-MN models described in the previous section, the construction of a complete motor nucleus requires defining how morphological parameters (*l*_*j*_, *d*_*j*_) and voltage-gated channel dynamics 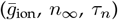 interpolate between the boundary units *α*-MN_1_ and *α*-MN_120_, ensuring population-level behavior consistent with recordings from the human FDI muscle [47]. This interpolation must ensure that the resulting motor pool faithfully reproduces both the orderly recruitment patterns associated with the size principle and the rate coding characteristics observed experimentally. This parameterization constitutes the first computational step in MyoGen’s EMG generation pipeline (Fig. 9B).

To accommodate diverse experimental conditions, we implemented two distinct motor pool parameterization models and their normalized version within a unified framework. This modular approach enables systematic comparative studies of recruitment patterns and rate coding behavior while maintaining flexibility for different applications. All models are implemented using literature-consistent notation that prioritizes computational stability. However, to enhance conceptual clarity in the presentation below, we reformulate the first model from its numerically robust implementation into a mathematically equivalent form that better conveys the underlying principles. Subsequent models, which extend this foundational approach, are presented directly in this more transparent notation.

The first model implements the classic formulation by Fuglevand et al. [25] for motor pool parametrization, in which the interpolation curve is defined by function in Eq. (33). In the original formulation, the parameter commonly denoted as the *recruitment range* represents the ratio between the highest and lowest MU recruitment thresholds. In our implementation, however, this parameter does *not* represent the physiological recruitment range; instead, we reinterpret it as an *exponential ratio* (ER) that purely controls the spread of the exponential interpolation. Accordingly, sf_F_[*i*] is the scaling factor of the *i*-th MU, and *N* is the total number of MUs in the motor pool.

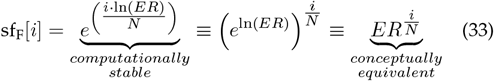

The model by De Luca and Contessa [26] incorporates a slope correction to allow for more physiological threshold distributions, as shown in Eq. (34), where *s* is the slope parameter, which allows the user to adjust the “elbow” of the distribution, effectively making it more or less linear.

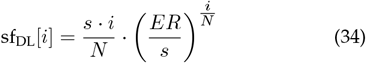

Konstantin et al. [4] introduces a normalized formulation of the Fuglevand et al. [25] model that provides explicit control over the maximum scaling factor. This normalization enables direct specification of the highest scaling value (sf_max_) while preserving the original motor-unit spacing of the Fuglevand et al. [25] model, as shown in Eq. (35).

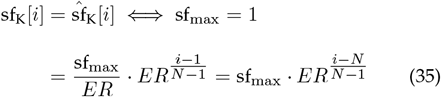

The final model is a normalized version of the De Luca and Contessa [26] formulation and is referred to in the API as the combined mode, as it integrates the key properties of the three preceding models (Eq. (37)). It preserves the shape characteristics of the De Luca model (Eq. (34)) while adopting the scaling strategy of Konstantin et al. [4] (Eq. (35)), enabling flexible fitting of recruitment patterns across muscles and pathological conditions.

The model is obtained by applying a min–max normalization (Eq. (36)) to the De Luca formulation, with bounds *a* = sfmax*/ER* and *b* = sfmax derived from Eq. (35), yielding the scaled factors shown in Eq. (37).

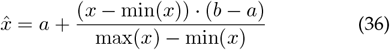

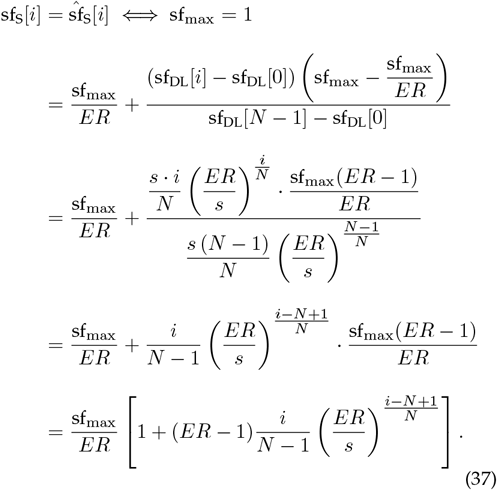

#### 5.3.6 Electrophysiological indicators

The electrophysiological properties (Fig. 5) of the computational models were estimated following the same procedure used in animal models [41, 42]. The parameters of the active properties of the model (Tab. 2) were adjusted to reproduce the electrophysiological indicators observed in experiments with anesthetized cats (*γ*_*CaL*_ = 0, passive dendrite), such as: the resting potential of the soma membrane (*V*_*rest*_); *R*_*in*_; *τ*_*m*_; Rheobase current (*I*_*rheo*_); AP amplitude; AHP amplitude and; AHP duration. With these electrophysiological indicators of the model adjusted to the experimental data, *γ*_*CaL*_ was defined as unitary, simulating the absence of anesthesia or the presence of neuromodulatory activity over the MN, and the parameters of *g*_*CaL*_ were adjusted to reproduce the hysteresis properties of the relation between the MN discharge rate and the injected current amplitude into the soma (*f* - *I*_*inj*_), observed in vivo in experiments with cats [43, 44].

##### 5.3.6.1 Relation between MN discharge rate and injected current

The simulation protocol to estimate the *F* - *I* relation (Fig. 5B & C) of the MN models adopted a somatic current injection with an ascending ramp profile with a slow rate of change (2.5 nA/s), approaching steady-state behavior [43]. The protocol consisted of a 1 s baseline period at 5 nA, followed by a 10 s linear ramp from 5 to 30 nA, and ending with a 1 s period at 30 nA. Spike times were detected when the membrane potential crossed 50 mV. Instantaneous DR was calculated from interspike intervals (ISIs), where each ISI was converted to frequency as *f* = 1000*/*ISI (with ISI in ms yielding frequency in Hz). To reduce variability, ISIs were smoothed using a 5-spike moving average, with the corresponding time point assigned to the center of the averaging window. For each time point, the injected current amplitude was determined from the ramp protocol timeline. The *f* - *I* gain was estimated as the slope of a linear fit (least-squares method) between the instantaneous discharge frequency and the injected current intensity, using only data from the ascending ramp phase (excluding the baseline and recovery periods).

##### 5.3.6.2 Rheobase current

The rheobase current *I*_rheo_ (Fig. 5D top left) estimates the excitation needed to activate a motor neuron, that is, the minimum current needed to generate an AP from a long-duration pulse injected into the soma. The simulation protocol used current pulses with increasing amplitudes of 0.1 nA and with a duration of 50 ms. An AP was detected when the somatic membrane potential exceeded the resting potential by at least 40 mV. The smallest pulse amplitude that promoted an AP was considered to be *I*_rheo_ [41].

##### 5.3.6.3 Membrane time constant

The experimental protocol to estimate the membrane time constant *τ*_*m*_ (Fig. 5D top right) consisted of the injection of rectangular pulses of somatic current with an amplitude of 20 nA and duration of 1 ms, applied after a 5 ms delay. The current injection hyperpolarizes the membrane potential briefly, and the interval from the peak hyperpolarization until the transmembrane voltage returned to within 0.1 mV of the resting potential was analyzed. A logarithmic transformation (ln |*V*_*s*_(*t*) −*V*_rest_|) was applied to the transmembrane potential during the recovery phase, and the line with the best fit to this curve was estimated using the least-squares method. The value of *τ*_*m*_ was estimated as the absolute value of the inverse slope of the fitted line [42].

##### 5.3.6.4 Input resistance

The experimental protocol to estimate the input resistance *R*_in_ (Fig. 5D middle left) consisted of injecting rectangular current pulses with a duration of 1000 ms and subthreshold amplitudes (between −5 and −1 nA) into the somatic compartment and measuring the steady-state membrane potential at the end of each pulse [41]. Then, *R*_in_ was estimated as the slope of the line with the best fit (least-squares method) to the points defined by the injected current intensity and the corresponding change in membrane potential from rest.

##### 5.3.6.5 Action potential and after-hyperpolarization (AHP)

The amplitude of the AHP (Fig. 5D middle right), as well as the duration of the AHP (Fig. 5D bottom row), were estimated from suprathreshold depolarizing current injections with a duration of 0.5 ms and a 5 ms delay [41]. The current amplitude was iteratively increased (starting at 35 nA, in 10 nA increments) until an AP with amplitude exceeding 40 mV above resting potential was elicited. The amplitude of the AP was measured as the difference between the peak transmembrane potential during the AP and *V*_rest_. AHP amplitude was measured as the difference between *V*_rest_ and the peak hyperpolarization observed immediately after the AP repolarization phase. AHP duration was estimated as the time interval between the AP peak and the moment when the membrane potential returned to within 0.15 mV of *V*_rest_ following the AHP. AHP half-decay duration was measured as the time interval from the peak hyperpolarization to the point where the membrane potential recovered to the midpoint between *V*_rest_ and the peak hyperpolarization (i.e., *V*_rest_ AHP amplitude*/*2).

##### 5.3.6.6 Parameter deviation

Parameter deviation D (Table S1) was defined as the normalized extent to which the simulated parameter range extended beyond the corresponding experimental range. Let *S*_min_ and *S*_max_ denote the minimum and maximum values of the simulated range, and *E*_min_ and *E*_max_ the minimum and maximum values reported in the literature. Deviations below and above the experimental range were quantified as max(0, *E*_min_ − *S*_min_) and max(0, *S*_max_ − *E*_max_), respectively. The total out-of-range extent was obtained by summing these contributions and normalizing by the width of the experimental range, (*E*_max_ −*E*_min_), yielding a percentage deviation (Eq. 38). The deviation evaluates to zero when the simulated range is fully contained within the experimental range.

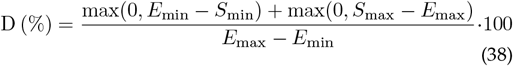

#### 5.3.7 Interneurons

Interneurons (INs) were modeled using a single-compartment configuration, following the same principles applied to *α*-MNs. Their morphological and electrotonic properties were tuned to match the experimental data of Bui et al. [54], assuming similar structural and electrophysiological characteristics for group II and Ib INs (gII and gIb). All INs shared the same set of voltage-gated ionic channels as the *α*-MNs and exhibited uniform dynamical behavior across the population. The IN models were previously validated against key biophysical properties, including frequency–current relations, afterhyperpolarization dynamics, and synaptic response profiles [5, 6].

The gII transmit signals with high efficacy, with EPSPs from a single group II afferent sufficient to evoke their discharge [55]. In contrast, during voluntary contraction of the agonist muscle, the Ib pathway reduces its transmission, an adaptive mechanism that prevents force-induced Ib inhibition from disrupting the sustained discharge of active *α*-MNs [56]. Renshaw cells and their recurrent inhibitory circuitry are predominantly associated with motor nuclei innervating proximal musculature and are largely absent or functionally negligible in distal limb muscles. Accordingly, recurrent inhibition was not implemented in the present model, which focuses on distal motor pools [57].

### 5.4 Dual-modality EMG simulation

Intramuscular and Surface EMG generation (i/s-EMG; Fig. 2 & Fig. 3) were implemented to allow cross-modal validation studies and comprehensive benchmarking of EMG processing algorithms.

sEMG simulations are based on the multilayer cylindrical volume conductor model by Farina et al. [58], which includes realistic anatomical tissue layers (skin, subcutaneous fat, muscle, and bone), each characterized by specific electrical conductivity and thickness. iEMG simulations follow the approach described by Konstantin et al. [4], using needle electrodes with configurable differential geometries and realistic insertion depths. This model accounts for motor-unit-specific detectability based on electrode proximity, allowing selective recording of individual muscle fibers.

From these two independent frameworks, we introduced several adaptations and improvements. The muscle fiber and MU territory distribution model from Konstantin et al. [4] was made compatible with the sEMG module, producing more physiologically realistic MUAP generation. The compatibility between the sEMG and iEMG modules enables direct cross-modal comparisons under identical physiological conditions.

The sEMG simulation in the original formulation of Farina et al. [58] relies on Bessel functions of order n, which can become numerically unstable for large n. To mitigate this, we adapted the simulation to compute the Bessel functions in logarithmic space using an exponential scaling approach, which preserves the correct values while avoiding overflow and underflow [59]. Additionally, the multi-loop computations were fully vectorized and implemented with parallel execution, enabling simultaneous evaluation across multiple elements and leveraging modern multi-core architectures to accelerate simulations. The handling of the electrode arrays was unified into a single interface that supports arbitrary spatial configurations for both surface and intramuscular electrodes.

### 5.5 Muscle-tendon unit and proprioceptive feedback models

MyoGen implements three interconnected biophysical models: a Hill-type muscle–tendon unit (MTU) model for force generation, a muscle spindle model for length/velocity sensing, and a Golgi tendon organ (GTO) model for force sensing. The muscle model outputs (force, length, velocity) drive the proprioceptive sensors for closed-loop sensorimotor control.

#### 5.5.1 Muscle unit

The mechanical event associated with the arrival of *α*-MN APs at the neuromuscular junctions is known as the twitch contraction. If the nonlinearities of the MU contraction process are neglected, this phenomenon can be approximated as the impulse response of a critically damped second-order system [25], whose temporal evolution is represented by the signal 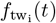. Accordingly, the MU force response 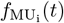 to a train of APs *e*_i_(*t*) can be approximated by the convolution between *f*_tw,i_(*t*) and the motor unit spike train (Equation 39).

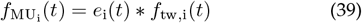

To improve computational efficiency, the force-generation process can be interpreted as linear and time-invariant (LTI) system, allowing the application of discretization techniques such as impulse invariance [60]. In Eq. 40, 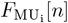 denotes the discrete-time version of the force generated by a MU (*f*_MU_(*n*)), Δ*t* is the sampling interval, *t*_p,i_ is the time to reach peak contraction, 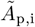 is the normalized contraction amplitude, and *e*_*i*_[*n*] is the discretized discharge train corresponding to *e*_*i*_(*t*).

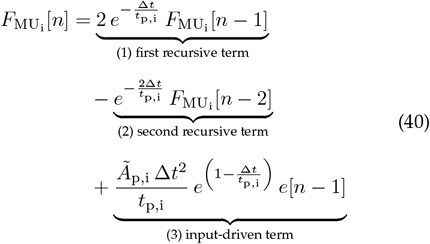

To model the nonlinear relation between MU force and *α*-MN discharge rate [61], reflecting the saturation dynamics of sarcomeric [Ca^2+^], we applied a sigmoidal scaling to the force output [13] (Eq. 41). The parameter *c* was iteratively determined using the secant method so that the first and last recruited MUs reached force saturation at 50 Hz and 100 Hz, respectively, corresponding to the physiological range reported for the human FDI muscle [62, 63].

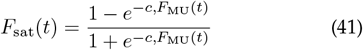

Motor units (MUs) within a muscle exhibit a wide range of contraction amplitudes (*A*_p,i_), spanning more than two orders of magnitude and scaling with *α*-MN size by approximately 130 times, ranging from 3 mN to 390 mN [62, 63]. Contraction time (*t*_p,i_) varies inversely with MU size so that early-recruited motor units contract more slowly and produce lower twitch forces, while later-recruited motor units contract rapidly and with a higher twitch force. Typical variations in contraction times range from two-to fivefold and were modeled here with a threefold ratio, ranging from 120 to 40 ms [25, 62, 64].

To convert MU force outputs into activation signals driving the contractile element of the muscle–tendon unit (MTU), we normalized the force generated by each MU by its tetanic force (*F*_tet,*i*_), defined as the force produced at saturation firing frequency. We then used the normalized activation signals (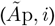, *i*) in Eq. 40 during the simulations. For simulations of force modulation driven by cortical beta-band input, we configured the motor pool to represent soleus musculature, comprising 800 type I and 100 type II motor units. For closed-loop reflex simulations with sensory feedback, we computed the activation signals *a*I(*t*) and *a*_II_(*t*) by summing the forces produced by the first 102 slow (type I) and the last 18 fast (type II) motor units, respectively, as defined in Eqs. 43 and 44 [63].

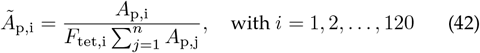

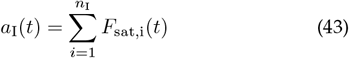

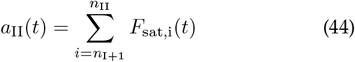

#### 5.5.2 Hill-Type muscle

Muscle force generation was modeled using a Hill-type formulation, following the classical framework by Zajac [65], which captures the nonlinear interactions between muscle fiber activation, length, and contraction velocity to produce physiologically realistic forces. The muscle–tendon unit (MTU) consists of a contractile element (CE), which receives the activation signals *a*_I_(*t*) and *a*_II_(*t*) from slow and fast *α*-MNs, in parallel with a viscoelastic element characterized by an elastic constant *K*_PE_ and a viscous constant *B*_PE_. A nonlinear series elastic element (*K*_SE_) represents the tendon and distal aponeurosis. The force generated by the CE is transmitted to the tendon considering a pennation angle (*α*) between the muscle fibers and the line of action of the tendon, such that only the component of the fiber force aligned with the tendon contributes to the MTU force.

The force generated by the active contractile element, 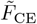(normalized by the unit activation force *F*_0_), is defined in Eq. 45 as a function of type I (*a*_I_) and type II (*a*_II_) motor unit activation levels, as well as the force–length (*f*_l_, Eq. 46) and force–velocity (*f*_v_, Eq. 47) relations of each type of muscle fibers. Only the activation levels (*a*_I_, *a*_II_), the normalized muscle length 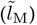, and the normalized muscle velocity 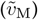 are functions of time, *t*; all other terms are treated as time-invariant parameters. For clarity, the explicit time dependence is omitted in the equations. Here, 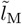 and 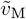 denote the muscle length and velocity normalized by the optimal fascicle length, 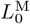. The MTU model parameter values are summarized in Tables 3 and 4.

**TABLE 3:**
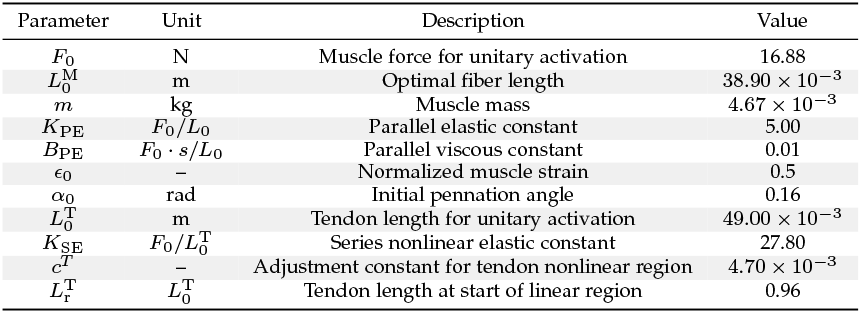
Parameter values for the muscle–tendon unit model corresponding to the human FDI muscle, used in the closed-loop reflex simulations [6, 66–68].

**TABLE 4:**
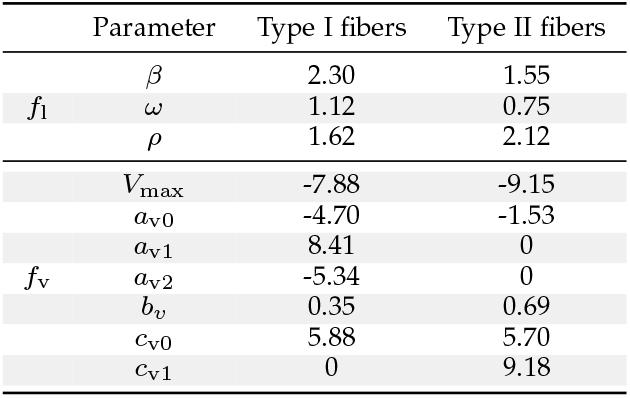
Parameter values for the force–length (*f*_l_) and force–velocity (*f*_v_) curves of type I and type II muscle fibers [6, 67, 68].

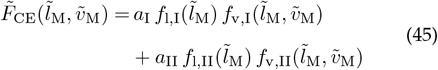

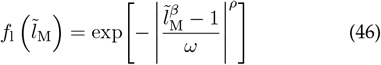

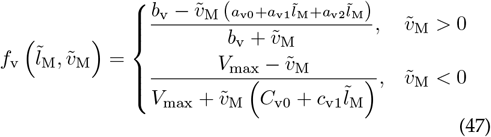

The force associated with the viscoelastic elements in parallel with the contractile element is defined by Eq. 48, where 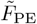 is the parallel viscoelastic force normalized by *F*_0_, *K*_PE_ is the element elastic constant, *B*_PE_ is the element viscous constant, and *ϵ*_0_ is the normalized muscle strain.

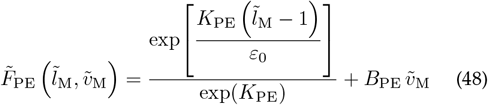

The force generated by the tendon, represented by the nonlinear series elastic element (*K*_SE_) connected in series with the muscle, was defined by Eq. 49. In this equation, 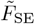 denotes the normalized tendon force with respect to the maximum isometric force *F*_0_, 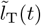 is the normalized tendon length relative to the tendon length at unit activation 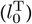, *K*_SE_ is the tendon stiffness constant, *c*_T_ is a shape parameter defining the nonlinear region, and 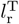 is the tendon length at which the linear region begins.

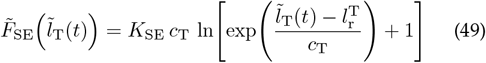

The tendon length (*l*_T_(*t*)) and muscle length (*l*_M_(*t*)) are related by Eq. 50, where *l*_MT_(*t*) denotes the total muscletendon unit (MTU) length and *α* represents the muscle fiber pennation angle. The pennation angle *α* depends on the normalized muscle length 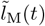 and the initial pennation angle *α*_0_ (see Eq. 51).

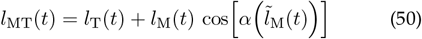

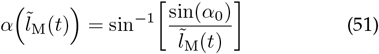

The *l*_MT_ of the FDI and its moment arm (*r*_FDI_) relative to the metacarpophalangeal joint were estimated from values obtained using a previously developed biomechanical model of the index finger [66] with the aid of OpenSim [69]. The estimation was performed by fitting polynomial curves that minimized the mean squared error relative to inverse kinematics data extracted from the biomechanical model [70]. Eqs. 52 and 53 describe the fitted curves as functions of the metacarpophalangeal joint angle (*θ*_*A*_), with positive angles corresponding to abduction. The polynomial coefficients are listed in Table 5, where *A*_*K*_ and *B*_*K*_ denote the coefficients for fitting the MTU length (*L*^*MT*^) and the moment (*m*), respectively.

**TABLE 5:** Polynomial fit parameters for the MTU length (*L*^MT^) and moment arm (*M*) of the FDI muscle (Eqs. 52 and 53).

| Parameter [unit] | Value |
| --- | --- |
| $A_0$ [m] | $8.52 \times 10^{-2}$ |
| $A_1$ [m/deg] | $-1.18 \times 10^{-4}$ |
| $A_2$ [m/deg<sup>2</sup>] | $-4.63 \times 10^{-7}$ |
| $A_3$ [m/deg<sup>3</sup>] | $9.42 \times 10^{-10}$ |
| $A_4$ [m/deg<sup>4</sup>] | $4.85 \times 10^{-12}$ |
| $B_0$ [m] | $6.83 \times 10^{-3}$ |
| $B_1$ [m/deg] | $4.84 \times 10^{-5}$ |
| $B_2$ [m/deg<sup>2</sup>] | $3.69 \times 10^{-8}$ |
| $B_3$ [m/deg<sup>3</sup>] | $6.31 \times 10^{-10}$ |
| $B_4$ [m/deg<sup>4</sup>] | $-6.36 \times 10^{-11}$ |

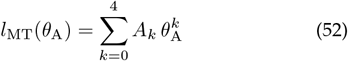

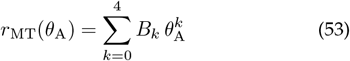

Applying Newton’s laws to the mechanical system representing the MTU yields Eq. 54, which is integrated in two steps to obtain the velocity (*v*_MT_) and length (*l*_MT_) of the MTU.

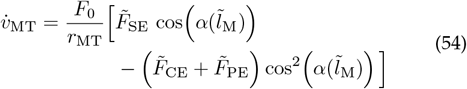

#### 5.5.3 Muscle Spindle

Muscle spindle dynamics were implemented according to the formulation of Mileusnic et al. [71]. The model was originally parameterized to optimally fit afferent recordings from the cat medial gastrocnemius muscle. Subsequently, we adjusted the gain parameters to more accurately reproduce the afferent activity observed in the human FDI muscle [72]. The receptor comprises three intrafusal muscle fiber types, each with distinct mechanical and activation properties: dynamic (*b*_1_) and static (*b*_2_) nuclear bag fibers, and nuclear chain (*c*) fibers [73]. The model reproduces the transduction of muscle length, velocity, and fusimotor drive into primary (Ia) and secondary (II) afferent discharge.

Dynamic (*γ*_dyn_) and static (*γ*_stat_) fusimotor drives modulate intrafusal activation through first-order low-pass dynamics. The activation levels of the dynamic and static bag fibers (*a*_b1_ and *a*_b2_, respectively) evolve according to the differential equations shown in Eqs. 55 and 56. The parameters *p, f*, and *τ* denote fiber-specific constants (see Table 1 in Mileusnic et al. [71]).

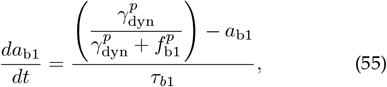

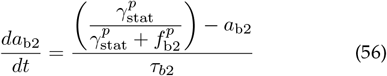

Each intrafusal muscle fiber was modeled as a coupled spring–mass–damper system representing two regions: a sensory region (*T*_SR_) and a polar region (*T*_PR_). The model incorporates nonlinear force–velocity relations and fusimotor-dependent modulation of stiffness and damping. The coupled differential equations governing *T* (*t*) and 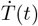 were solved numerically using a fourth-order Runge–Kutta integration method.

Afferent responses were computed as rectified functions of intrafusal stretch. The primary (Ia) and secondary (II) afferent discharge rates were obtained from the expressions shown in Eqs. 57–58. The Ia gains (*G*_*i*_) were set to 6500 for bag_1_ fibers and 3250 for both bag_2_ and chain fibers, whereas for group II afferents *G*_*i*_ was set to 3500 for both bag_2_ and chain fibers.

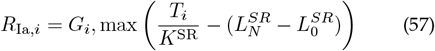

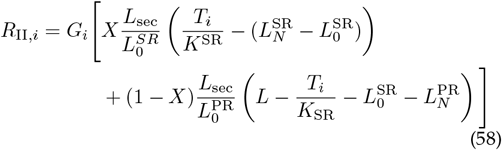

Primary afferent output was computed through the occlusion mechanism described in Eq. 59, and the secondary output as the sum *R*_II_ = *R*_*b*2_ + *R*_*c*_.

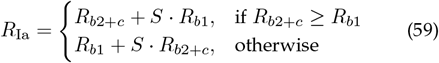

The overall process can be summarized as a cascade transforming the muscle kinematics into afferent mean discharge rates:

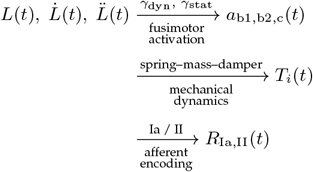

where *L*(*t*), 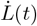, and 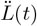 denote the instantaneous muscle length, velocity, and acceleration, respectively; *γ*_dyn_ and *γ*_stat_ represent the dynamic and static fusimotor activations driving intrafusal fibers; *a*_b1,b2,c_(*t*) are the corresponding intrafusal activations for bag_1_, bag_2_, and chain fibers; *T*_*i*_(*t*) is the resulting intrafusal tension; and *R*_Ia,II_(*t*) are the primary (Ia) and secondary (II) afferent discharge responses.

#### 5.5.4 Golgi tendon organ

The GTO was modeled as a non-linear force sensor, so that a logarithmic function was employed to provide the saturation of Ib DR with the increase of force, with a linear dynamics, following the formulation of Lin and Crago [74]. Extrafusal muscle force *F*_*m,e*_(*t*) was transduced into an afferent drive using the formulation in Eq. 60. The gain and scaling constants were calibrated to human data (*G*_*g*_ = 40 Hz, *G*_*f*_ = 4 N) from Aniss et al. [75].

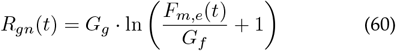

The GTO dynamics were reformulated as a second-order digital filter via bilinear transformation, ensuring numerical stability and temporal fidelity in discrete time [6]. The resulting discrete-time calculation is given in Eq. 61, where the coefficients *b*_0_, *b*_1_, *b*_2_, *a*_1_, *a*_2_ were derived from the continuous-time transfer function. A final half-wave rectification enforced nonnegative DR, as shown in Eq. 62.

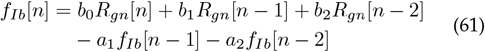

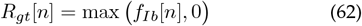

The overall processing cascade can be summarized as:

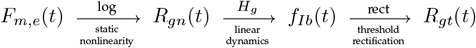

#### 5.5.5 Afferents Ia, II, and Ib

We modeled muscle receptor outputs to represent the mean discharge rates of Ia and II afferents from muscle spindles and Ib afferents from Golgi tendon organs. These outputs drove independent, non-homogeneous Gamma renewal point processes for each afferent population [37] (see section 5.2). We adjusted the shape factors of the point processes to reproduce the ISI coefficient of variation, setting 8.3% for group I afferents (Ia and Ib) and 3.6% for group II [76].

We assigned the total number of afferents based on literature estimates [77, 78], including 73 Ia afferents, 80 group II afferents (110% of the Ia count), and 58 Ib afferents (80% of the Ia count). Conduction velocities were set according to values reported for distal upper-limb muscles: approximately 68 m · s^−1^ for Ia and Ib afferents [56, 79], and about 40 m · s^−1^ for group II afferents [80]. Ia afferents formed direct, excitatory monosynaptic projections to homonymous *α*-MNs, representing the canonical monosynaptic reflex arc, with a connection probability of 80% that reflects their homogeneous excitatory distribution and strong influence to the motor pool. In contrast, Ib and II afferents connected to INs with a probability of 30%, capturing their predominantly indirect, di-synaptic influence within spinal circuitry [6, 56].

Each sensory afferent unit *i* within a population *X* ∈ Ia, Ib, II began discharge when the output of the corresponding muscle receptor model 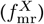 exceeded its recruitment threshold (*rt*_*X,i*_), which was linearly distributed across the population between 0 and 40 Hz to reproduce the recruitment behavior of afferent fibers observed in human experiments [81]. The mean DR of each afferent (*f*_*X,i*_) was then computed according to Eq. 63, where *ifr*_*X,i*_ denotes the initial DR, randomly varied by a Gaussian random variable 𝒩 (5, 2.5^2^) to introduce physiological variability among units.

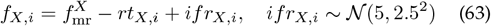

### 5.6 Finetuning pipeline for MUs

To fine-tune the MU pools (Fig. 4), we implemented a three-step optimization pipeline.

In the first step, we determined the parameters required for each cortical pool to drive a 100-MU pool at a target DR. Cortical pools were modeled as gamma point processes with a predefined shape parameter, and this step estimated the input characteristics necessary to reproduce the desired MN discharge statistics.

We optimized the following parameters using Optuna [24], a hyperparameter optimization library for Python:

- number of cortical neurons (range: 100–1000)
- connection probability between a cortical neuron and an MN (range: 0.0–1.0)
- mean spiking rate of cortical neurons (range: 10– 250 Hz)

All parameters were optimized using a Tree-structured Parzen Estimator (TPE) [82]. At each iteration, TPE constructed two Gaussian mixture models: *l*(*x*), representing parameter values from the best-performing trials, and *g*(*x*), representing all others. New candidate parameters were sampled to maximize the expected improvement, quantified as the ratio *l*(*x*)*/g*(*x*).

The optimization minimized the following cost-function:

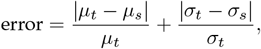

where *µ*_*t*_ and *µ*_*s*_ denote the means of the target and simulated DR distributions, respectively, and *σ*_*t*_ and *σ*_*s*_ their corresponding standard deviations. The DR standard deviation was computed following Eq. 9 in Watanabe et al. [13].

In the second step, we computed the resulting muscle force using the model described by Fuglevand et al. [25], assuming that the output corresponded to 100% of the MVC. Finally, in the third step, we optimized the cortical spiking rate identified in step one to produce muscle forces corresponding to specified submaximal contractions (as a percentage of MVC).

To quantitatively evaluate how well the simulated MU discharge statistics reproduced the experimental data distribution, we introduced an asymmetric geometric coverage metric (Fig. 4B). This metric specifically addressed the question: *What fraction of the experimental distribution can be reproduced by the simulator?*

For each variable, we estimated probability density functions from both experimental and simulated datasets using Gaussian kernel density estimation (KDE). Each KDE was normalized to emphasize geometric similarity rather than absolute probability mass, thereby preventing bias from unequal sample sizes between datasets.

The geometric coverage *C* (in %) was defined as Eq. (64).

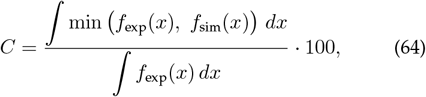

where *f*_exp_(*x*) and *f*_sim_(*x*) denote the normalized experimental and simulated density functions, respectively, evaluated over the physiologically relevant domain *x*. The numerator quantifies the intersection area between the two distributions via the point-wise minimum function, while the denominator represents the total area under the experimental distribution. Integrations were performed numerically using the trapezoidal rule with 500 evenly spaced evaluation points across the domain.

The resulting metric ranged from 0% (no overlap) to 100% (complete coverage of the experimental distribution by the simulation). Importantly, the metric is asymmetric: high coverage indicates comprehensive simulation performance relative to experimental data, but does not penalize exploration of regions beyond those sampled experimentally.

### 5.7 Spike train agreement analysis

Decomposition accuracy (Fig. 3G) was assessed by comparing algorithmically identified spike trains to corresponding ground-truth discharges using a two-stage framework: temporal spike matching followed by optimal MU assignment.

#### 5.7.1 Temporal spike matching

Spikes were matched within a ± 5.0 ms tolerance window for the iEMG signals and ± 12.5 ms for sEMG. A decomposed spike was labeled a true positive (TP) if it occurred within the tolerance window of a ground-truth spike. Greedy nearest-neighbor matching enforced one-to-one correspondence, preventing multiple assignments of the same ground-truth event. Unmatched spikes were classified as false positives (FP) or false negatives (FN).

#### 5.7.2 Optimal motor unit assignment

Because MU identifiers assigned by decomposition algorithms are arbitrary, correspondence between ground-truth and decomposed units was resolved by constructing an *N* ×*M* agreement matrix containing the F1 scores for all unit pairs. For each matched pair, sensitivity (recall) was computed as TP*/*(TP + FN), precision as TP*/*(TP + FP), and the F1 score as

2 · (precision · sensitivity)*/*(precision + sensitivity).

The Hungarian algorithm [83] was applied to find the one-to-one mapping that maximized the total F1 score. Unmatched units were reported as either missed detections (ground truth) or false discoveries (decomposed motor units).

### 5.8 EMG processing for dimensionality reduction

The EMG data produced in Fig. 8B & H was simulated at 2048 Hz from two 8 × 8 electrode grids (128 channels total: 64 agonist, 64 antagonist) and preprocessed using the following pipeline. Signals were first bandpass filtered using a 4th-order Butterworth bandpass filter from 20–500 Hz, with the filter applied bidirectionally (zero-phase) to prevent temporal distortion. A Longitudinal Single Differential (LSD) spatial filter was then applied independently to each electrode grid to enhance local muscle activity and suppress common-mode noise. Spatially filtered signals were full-wave rectified to extract the amplitude envelope, then segmented into overlapping windows of 320 samples (≈150 ms) with a shift of 8 samples (≈4 ms), yielding approximately 96.9% overlap between consecutive windows. Each channel was normalized to the range [0, 1] using minmax scaling, where global minimum and maximum values were computed across all signal-to-noise conditions to ensure comparable scaling. Prior to dimensionality reduction, features were centered and scaled using robust statistics (median and interquartile range) to minimize the influence of outliers and balance channel contributions.

Dimensionality reduction was performed using principal component analysis (PCA) from the `scikit-learn` library and Uniform Manifold Approximation and Projection (UMAP) from the GPU-accelerated `cuML` implementation (RAPIDS AI; [84]), enabling efficient processing of high-dimensional EMG data. For UMAP, we set 125 neighbors, a minimum distance of 0.5, and 500 training epochs, while all other parameters were kept at their default values in cuML version 25.12.00.

## Data and Code Availability

The MyoGen simulation framework, analysis scripts, and example datasets are publicly available on GitHub (https://github.com/NsquaredLab/MyoGen) and Zenodo (DOI: 10.5281/zenodo.18078175). Documentation is available at https://nsquaredlab.github.io/MyoGen/.

## Acknowledgment

We are grateful to our colleague Finja Beermann for her help and insights on neurophysiological questions and for her help with the early version of this manuscript.

Parts of Figs. 1, 3A, 4A, and 7A were created in https://BioRender.com, which we gratefully acknowledge..

## Supplementary

**TABLE S1:** Comparison of simulation parameters with literature values [41]. Parameters are sorted by order in which they appear in Zengel et al. [41]. Deviation denotes the percentage by which the simulated range exceeds the experimental range; wr (within range) indicates that the simulated values are fully contained within the literature’s range.

| Parameter | Ours | Literature | Deviation |
| --- | --- | --- | --- |
| Input Resistance ($M\Omega$) | 0.5 – 1.8 | 0.1 – 3.0 | wr |
| Memb. Time Const. (ms) | 3.0 – 11.2 | 3.0 – 15.0 | wr |
| AHP Magnitude (mV) | 4.0 – 4.8 | 1.0 – 10.0 | wr |
| AHP Duration (ms) | 34.8 – 136.0 | 30.0 – 270.0 | wr |
| AHP Half-Decay Duration (ms) | 10.7 – 42.6 | 12.0 – 60.0 | 2.7% |
| Rheobase Current (nA) | 5.3 – 18.9 | 4.0 – 40.0 | wr |

## Notes

This work was supported by the Bavarian Ministry of Economic Affairs, Regional Development and Energy (StMWi) under grant MV-2303-0006, by the European Research Council (ERC) through the project GRASPAGAIN under grant 101118089, and by the German Federal Ministry of Education and Research (BMBF) through the project MYOREHAB under grant 01DN23002 to ADV; the Coordenação de Aperfeiçoamento de Pessoal de Nível Superior - Brasil (CAPES) - Finance Code 001 to RNW; the National Council of Technological and Scientific Development (CNPq) under grant number 147239/2024-9 to RLB; the Brazilian Public Ministry of Work (MPT) under grant 002118.2019 to RGM; the National Council of Technological and Scientific Development (CNPq) under grant number 421810/2023-8 to LAE, who is also a Research Productivity Fellow of CNPq (process number 316320/2023-4).

https://github.com/NsquaredLab/MyoGen

https://doi.org/10.5281/zenodo.18078175

### Competing Interest Statement

The authors have declared no competing interest.

